# Expanding the fly eye gene regulatory network: from *Drosophila* to the hoverfly *Episyrphus balteatus*

**DOI:** 10.1101/2025.09.02.673659

**Authors:** T. Navarro, J. Torres, R. Sáez-Moreno, G. Guerrero, I. Almudí, A. Iannini, J. Figueras, K Wotton, S. Aerts, F. Casares

## Abstract

Comparing developmental mechanisms across long evolutionary distances is challenging, especially when the level of description in the compared species is very disparate. This is the case for the gene regulatory network (GRN) that underlies the development of compound eyes in insects, where the very detailed level description of the network in the fruit fly *Drosophila melanogaster* contrasts with the knowledge of the network in other well studied insects, such as the flour beetle *Tribolium* or the desert locust *Schistocerca.* It is not even known to what extent the *Drosophila* eye GRN is typical of flies (diptera). Here we have introduced the marmalade fly *Episyrphus balteatus* (Syrphidae), separated from *Drosophila* approximately 90 MYrs, as a fly species to compare the mechanisms of compound eye development. The *Episyrphus* eye develops from an imaginal disc which is, in the overall, similar to that of *Drosophila.* By generating parallel data sets in *Episyrphus* and *Drosophila* to maximize comparability, our results show how the genes known to participate in the early specification of the *Drosophila* eye and the onset of its differentiation are likely participating in the eye GRN of *Episyrphus,* further expanding the set of conserved genes/nodes in this network. This gene set might be the basis for further explorations of the developmental mechanisms involved in compound eye development beyond flies. By combining RNAseq and ATACseq profiling with TF motifs we derive the first eye GRN in *Episyrphus.* When built using *Episyrphus/Drosophila* conserved nodes, the resulting GRN model exhibited abundant internal connectivity, suggesting that, as proposed in *Drosophila,* the GRN is knitted with many regulatory feedbacks. Using link conservation as a criterion for discovering new regulatory interactions, we find that the AML1/Runx transcription factor *lozenge (lz)* is a negative regulator of the retinal determination gene *dachshund (dac)* in *Drosophila.* However, although many of the predicted regulatory links are conserved, a large number of regulatory connections seem species-specific, even among conserved nodes. This suggests quite a significant regulatory rewiring during the 90Myr period separating *Episyrphus* and *Drosophila*.

## INTRODUCTION

The development of eyes across animals, and despite their very different architectures, requires a basic genetic tool kit, composed of a set of conserved transcription factors (TFs) and signaling molecules. This fact suggests that all animal eyes may share a deep developmental homology (Erclik et al., 2009). However, the development of the different eye types, being these single lens camera type or compound (comprising many individual eyes), is also remarkably different. In order to understand to what extent the processes governed by the set of conserved TFs and signaling pathways are conserved or divergent, two further levels of description are required. First, one requires knowing which genes lie downstream of these conserved elements; and second, how these factors are integrated into a gene regulatory network-level description. A comprehensive description of eye development at this level of resolution is lacking for most eye types. An exception is the *Drosophila melanogaster* eye. *Drosophila* is a minute fly whose eye has been used as paradigm of compound eyes, the most abundant eye type in nature. Developmental genetics studies have already uncovered many of the genes essential for eye development and established some of their mutual cross-regulations. This work was initially complemented with the analysis of *cis*-regulatory elements (mostly activating elements called “enhancers”) to establish the skeleton of an eye GRN (the so-called Retinal Determination Gene regulatory Network, RDGN (Kumar, 2009)). Further research, including transcriptomics, epigenomics and genome-wide TF binding, and more recently single-cell omics, has resulted in more complete eye GRN models where space (that is, a model of the actual neuroepithelium) is included (Bravo Gonzalez-Blas et al., 2020; Potier et al., 2014; Sanchez-Aragon et al., 2019). The ultimate test for these models would be the correct prediction of eye phenotypic changes when specific elements in the GRN are perturbed, something that is still pending. Globally considered, the *Drosophila* eye is one of the most highly-developed models of organ specification and differentiation (Casares and Almudi, 2016).

The eye GRN is deployed in the posterior region of the eye-antennal imaginal disc, which gives rise to most structures of the adult head. The first action of the GRN is the specification of this posterior region as the eye progenitor domain. Then, signals derived from the disc’s posterior margin, mostly the *hedgehog* morphogen (Hh), start the recruitment of eye progenitors into retinal precursors first, which then undergo further differentiation. As these early photoreceptors also express Hh, the differentiation of the retina proceeds like a wave, with differentiating photoreceptors recruiting progenitor cells via Hh signaling cells being recruited anterior to the wavefront to initiate their differentiation (Treisman, 2013). The differentiation wavefront is characterized by an indentation in the disc’s epithelium, called the Morphogenetic Furrow (MF), and is used as a morphological reference of the differentiation process. Posterior to the MF, differentiating cells either exit the cell cycle or undergo a last round of mitosis before differentiating. The products of this differentiation process are clusters of photoreceptor cells, surrounded by lens-secreting and pigment-producing cells. Each of these cartridges is a unit eye, or ommatidium, which is identifiable at the eye surface by its covering lens (or facet). During this whole process, the action of Hh and other intercellular signaling molecules animate the GRN to define the developmental trajectories going from progenitors to the different cell types forming the ommatidium (Casares and Almudi, 2016; Treisman, 2013).

Even within insects, the level of resolution of the eye GRN is far from that of *Drosophila* (Bushbeck and Friedrich, 2008; Friedrich, 2006; Kittelmann and McGregor, 2024). Work in beetles (*Tribolium*; (Yang et al., 2009a; Yang et al., 2009b), honeybees (*Apis*; (Hu et al., 2024), crickets (*Grillus,* (Takagi et al., 2012)) and grasshoppers (*Schistocerca,* (Dong and Friedrich, 2010)), among others, has focused on specific elements of the GRN, finding out that the genes known to be important for eye development in *Drosophila* are generally also relevant for eye development in these species, although sometimes the discovered function does not fully correspond to that previously identified in *Drosophila* (see, for example, the analysis of the role of the *eyg* TF in *Tribolium* eye development (ZarinKamar et al., 2011)). This is not totally unexpected, because *Drosophila* development is quite derived within insects. While in most insects eyes develop progressively in the cephalic region of the larval ectoderm (hemimetabolous development), in flies it takes place internally, as an imaginal disc, and happens relatively rapidly, spanning the last phase of the larval life ((Ready, 1989), holometabolous development). The sparsity of the phylogenetic sampling means that we don’t even know whether eye development in *Drosophila* is typical within Diptera, an order where eyes exhibit a great morphological and functional diversification (Casares and McGregor, 2020). To tackle this issue, we have developed the first eye GRN model for the hoverfly *Episyrphus balteatus* and compared it to that of *Drosophila melanogaster*.

The evolutionary divergence between *Episyrphus* and *Drosophila* is approximately 90 million years (Myrs), placing it between the Mediterranean fruit fly, *Ceratitis capitata* (140 Myrs), and the tsetse fly, *Glossina morsitans* (65 Myrs) (http://www.timetree.org/). *Episyrphus balteatus* (De Geer, 1776; Diptera: Syrphidae), commonly known as the marmalade hoverfly, is a cosmopolitan species widespread across the Palearctic region. It plays important ecological roles, both through its aphidophagous larval stage and as a generalist pollinator during its adult stage. The adult *Episyrphus* has large compound eyes. This large size may be related to the flying dexterity of this group of flies, capable of hovering, airborne mating and long-distance migrations (Omkar and Mishra, 2016; Wotton et al., 2019). In addition, annotated high-quality genome versions have been published (Doyle et al., 2022; Hawkes and Sivell, 2023). The *Episyrphus* eye GRN and its comparison with that of *Drosophila* should shed light on a number of still unresolved issues. First, what is the minimal set of genes to be part of the GRN. Second, whether the set of TFs involved in the specification, growth and early differentiation of the *Drosophila* eye is conserved between two distantly related fly species, and whether this set can be expanded. Third, whether the regulatory code controlling the expression of shared genes is conserved or not –that is, what is the degree of “rewiring” within the regulatory network. Finally, whether conserved regulatory links are predictive of *functional* regulatory interactions, at least when tested in *Drosophila*.

Here we show, through the comparative analysis of the eye transcriptomes, that very similar functions and processes are activated during eye development in both species, which share a set of genes that can be used as a reference for the further characterization of eye development in other species. We then connected the transcriptome as a GRN by linking expressed TFs to target genes through Differentially Accessible Regions (eye DARs, relative to wing chromatin accessibility) in the eye disc, derived from ATAC-seq, which very often function as activating <u>c</u>is-regulatory <u>e</u>lements (CREs) or *enhancers*. In order to validate the regulatory predictions of our GRN, we tested a predicted link between two key nodes, *lozenge (lz)* and *dachshund (dac)*, using functional assays in *Drosophila,* and showed that *lz* acts as a *dac* repressor, lending support to our GRN model. The global view of the resulting GRN models identified shared regulatory links, but also links predicted to be species-specific. The core GRN comprises 106 conserved genes, and includes a set of 22 TFs (“kernel”), which according to the GRN model, are heavily interconnected. The degree to which connectivity was conserved, though, varied across genes. When this connectivity was analyzed not as a whole, but discriminating by enhancers, again some gene orthologs showed more regulatory similarity than others.

Although our *Episyrphus/Drosophila* conserved GRN model is surely incomplete, among other things because we still lack DNA binding motifs for a few conserved TFs, it represents the largest expansion of the eye GRN from *Drosophila* to another dipteran beyond drosophilids. In this way, this GRN could be considered a tentative minimal eye GRN model within dipterans, to be used as a reference for comparative studies in other Dipterans and beyond.

## RESULTS

### *Episyrphus* eye development resembles that of *Drosophila*

The eyes of *Episyrphus* are very large compound eyes. Compared with the Drosophila eye, composed by about 800 ommatidia, the *Episyrphus* eye comprises more than 3500 (Figure 1A,B). In addition, the eyes are sexually dimorphic, with the male eyes extending all the way to the dorsal midline (a trait called “holoptic eyes”), through this dimorphism was not analyzed further. In order to start investigating *Episyrphus* eye development we dissected L3 larvae at different time points and fixed the eye imaginal discs (Figure 1C). The eye disc is kidney-shaped, with the concave region pointing towards posterior. A furrow, similar to the morphogenetic furrow in *Drosophila* is detected, separating an anterior to a posterior region. This latter shows clustering of cells into ommatidia. As development proceeds, the area behind the MF increases, indicating a wave-like progression of the differentiation process. In addition to abundant mitotic cells anterior to the MF, as expected for progenitor cells, we also observe, especially midway through development, mitotic cells in the differentiating area, not particularly concentrated after the MF -as it is the case in *Drosophila* in the so-called second mitotic wave (Wolff and Ready, 1991). Overall, eye development in *Episyrphus* is similar to that in *Drosophila,* progressing in a wave-like fashion across the growing disc epithelium, although some aspects, such as the curvature of the MF or the pattern of proliferation behind it differ. Next, we describe the steps towards the generation of its GRN.

**Figure 1.**
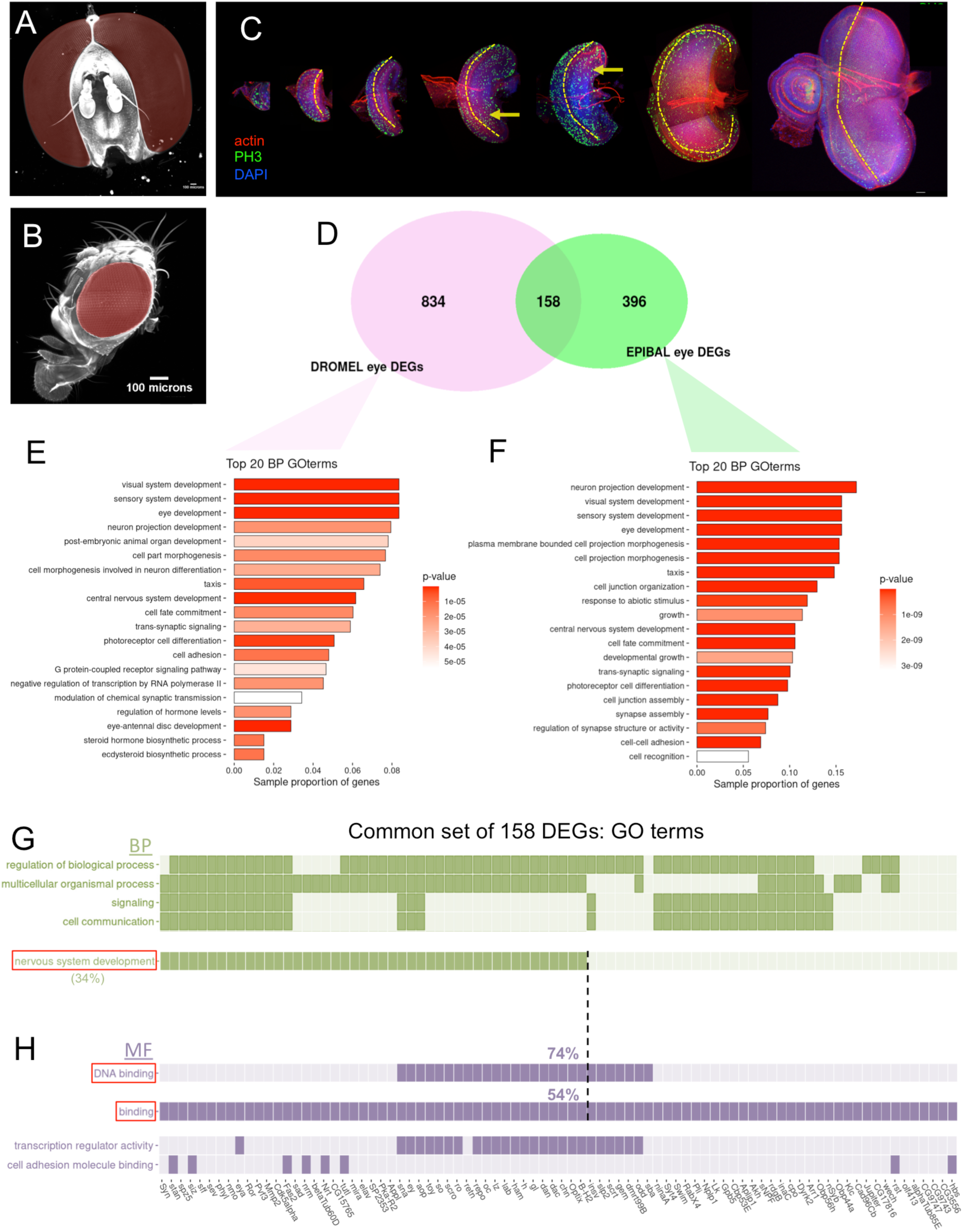
The *Episyrphus* and *Drosophila* eyes and their comparative transcriptomics. (A,B) Light sheet images of the adult heads of *Episyrphus balteatus* (A, male) and *Drosophila melanogaster* (B) to the same scale (scale bar = 100 micrometers). Eyes are pseudocolored in red. (C) *Episyrphus* eye discs at progressive stages of their development, stained for actin (rhodamine-phalloidin), DNA (DAPI) and the mitotic marker phosphorylated Histone H3 (PH3). The morphogenetic furrow (MF, marked by the dashed yellow lines) is not straight, but convex. Mitotic cells are abundant anterior to the MF. In addition, frequent mitotic cells are detected scattered throughout the differentiating region of the disc, posterior to the MF (indicated by the yellow arrows). (D) Venn diagram indicating the number of differentially expressed genes between eye and wing discs (“eye DEGs”) in *Drosophila* (“DROMEL”, pink) and *Episyrphus* (“EPIBAL, green). 158 eye DEGs are shared between the two species. (E, F) Top 20 Biological Process (BP) gene ontology terms for the *Drosophila* (E) and *Episyrphus* (F) eye DEGs. (G, H) eye DEGs associated with the top five GO terms in the Biological Process (G) or Molecular Function (H, where only 4 terms were enriched) among the shared 158 eye DEGs. A dark cell indicates that a gene (column) is associated with the GO term in the respective row. 34% of the genes belong to the BP term *nervous system development*. Within the set of genes associated with the MF term *binding*, the proportion of genes associated with *nervous system development* rises to 54%, further increasing to 74% when considering only those associated with *DNA-binding*.

### Identification of the nodes in the eye GRN

In a GRN model, the expression of any gene in the network is the result of the (activating or repressing) regulatory links connecting the gene’s *cis-*regulatory elements (CREs) to its upstream TF regulators. CREs (most common CREs are activating or *enhancer* elements) can be identified as chromatin regions with increased biochemical accessibility relative to the surrounding chromatin as assessed by a number of techniques (Arora and Tollefsbol, 2021), including <u>a</u>ssay for transposase-<u>a</u>ccessible <u>c</u>hromatin using sequencing (ATAC-seq (Buenrostro et al., 2013)). These regions are thereof “<u>d</u>ifferentially <u>a</u>ccessible regions” or DARs. A link between TF-A and a gene *B* is predicted to exist if A and *B* are co-expressed and there is a DAR in the vicinity of *B* containing a transcription factor binding site for A. Then, B is said to be a putative regulatory “target” of TF-A. These predicted links could be further experimentally validated by biochemical and genetic techniques. Therefore, as a first approximation, three types of data must be gathered in order to knit a GRN model: gene transcription (expression), DARs and TF binding sites. To build a first eye GRN model for *Episyrphus*, we obtained bulk RNA-seq and ATAC-seq data for eye and wing discs from mid-third instar larvae. To derive a comparable eye GRN in *Drosophila*, equivalent sets of data were obtained in this fly (see Materials and Methods/ RNA Sequencing and Bioinformatic Processing).

The first step in our GRN building strategy was to identify the sets of genes specifically involved in eye development. By comparing “eye” against “wing” discs we obtained the set of differentially expressed genes (DEGs) in the eye-antennal imaginal disc (*Episyrphus* and *Drosophila* eyeDEGs). This first gene set of eyeDEGs served to check whether the eye minus wing data sets captured eye development-specific processes. *Drosophila* and *Episyrphus* eye discs have 992 and 554 eyeDEGs, respectively (Figure 1D; Supp. Table 1). Gene ontology (GO) analysis of these eyeDEGs identified very similar overrepresented terms in both flies, including “visual system/sensory system/eye development” and “neuron projection development”, in agreement with the major cell populations of the eye discs at the stage studied: eye progenitors and early differentiating photoreceptor neurons (Figure 1E,F). The overlap between the two sets is of 158 genes (Figure 1D). This gene set is further enriched (relative to the total of eye DEGs) in terms such as “signaling” and “cell communication”, and molecularly in “binding”/”DNA binding” (Figure 1G,H), which includes transcription factors. Within both sets, the enrichment of genes belonging to the *nervous system development* term is even greater (Figure 1G,H), indicating that a key part of the common (conserved) DEGs set is composed of genes that regulate differentiation process.

The involvement of a gene in the eye GRN can be inferred, even in the absence of significant differential expression, if it is associated with regulatory regions that are specifically active in the eye —that is, eye DARs. One example helps clarify this point. The T-box *bicaudal* (*bi/omb)* gene is known to be expressed in both *Drosophila* eye and wing discs and to be required for the development of some of their adult derivatives (del Alamo Rodriguez et al., 2004; Tsai et al., 2015). However, our RNA-seq analysis did not identify *bi* as an “eye DEG”. And yet, when the ATAC-seq analysis was performed, the *bi* locus showed eye and wing-specific DARs, suggesting that different regulatory inputs are controlling *bi* transcription in these two primordia (Figure S1). Considering this, we decided to build the eye GRN with all the genes expressed in the eye (even if not *differentially* so), which are linked to one or more eye-specific DAR. DARs were linked to target genes using a proximity criterion to the nearest expressed gene (see Materials and Methods/DARs annotation). We called these genes “eye GDARs” (eye-expressed genes with linked eye-specific DARs) and became the nodes of our network models.

We detected 533 eye GDARs in *Episyrphus* which were GO-characterized, similarly to the eye DEGs, by terms such as “eye/visual/sensory system development”, “neuron projection development” and “photoreceptor cell differentiation”. In addition, this gene set is characterized by terms such as “wing disc development” that may point to those genes which, in addition to their eye expression, have known functions in the development of this organ (as is the case for *bi/omb,* described above), or “head development” in agreement with the fact that the eye disc also contributes to the development of other head structures, including the head capsule and the antenna (Figure S3A, right). The 624 *Drosophila* eye GDARs were also characterized by the same terms as the *Episyrphus* equivalent set, although some additional general terms pertaining to development, such as “epithelial tube morphogenesis” and “post-embryonic animal organ development”, appeared as more significant (Figure S3B, right). However, and in general, the set of eye-expressed genes linked to eye-specific DARs (eye GDARs) pointed to very similar biological functions in both species. This similarity in GO-characterization extended further when only eye GDARs of eye DEGs were considered, with “visual system/sensory system/eye development” being the top terms (Figure S3A,B; left). This GO enrichment analysis supports the notion that GDARs can be considered the nodes of the GRN models.

### A core of 106 eye genes

We identified 533 and 624 eye GDARs in *Drosophila* and *Episyrphus*, respectively. Of these, 106 genes (which we call “core” genes) are shared between the two species. This core gene set showed GO enrichment in terms such as “visual system development” or “photoreceptor cell differentiation” relative to the union of 1157 *Episyrphus* and *Drosophila* eye GDARs (Figure 2A-C). This was remarkable taking into account that the *Episyrphus* and *Drosophila* eye GDAR sets were already significantly enriched in these terms. We checked the expression of these genes in the single cell atlas dataset generated by Bravo Gonzalez-Blas and co-workers. (Bravo Gonzalez-Blas et al., 2020). All genes showed expression in some of the cell types of the eye disc epithelium but for *stumps,* which showed expression in glial cells. Among the core gene set, 29 genes have been associated to one or several signaling pathways, including the Hh, wg/Wnt TGF-β, with components of the MAPK pathway being particularly enriched (Table 1).

**Figure 2.**
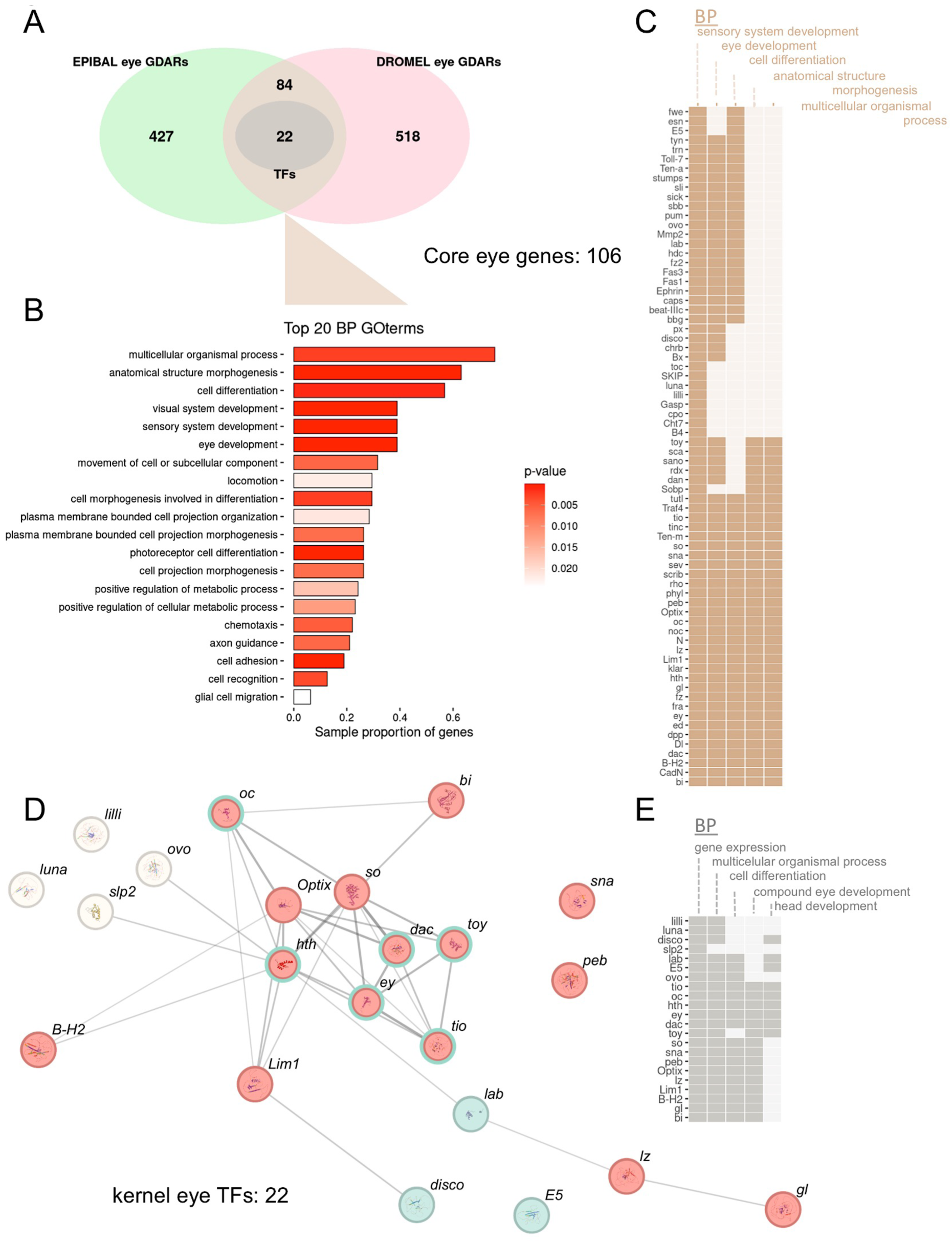
Analysis of the *Episyrphus* and *Drosophila* eye GDARs. (A) Venn diagram of the intersections between the two gene sets. Among the shared 106 genes (“core” gene set), there are 22 common TFs, the “kernel” TF set. (B) GO analysis of the core 106 gene set, showing the 20 top Biological Process (BP) terms. (C) List of genes, among the 106 set, with annotated terms from the 5 top GO terms. Dark cells indicates that a gene (row) is associated with the GO term in the respective column. (D) STRING interaction networks of the TF kernel set. Increasing link thickness indicates higher confidence. In red, the TFs associated with eye development; in cyan, the TFs associated with head development (TFs associated with both processes share both colours); and in white, the TFs not associated with neither of the two processes. The network is highly connected. (E) Kernel TF set and the assignation of the relevant (non redundant) five GO terms to each of them. As above, dark cells indicates annotation of the gene (row) with the specific GO term (columns). Note that the automatic GO annotation of *lilli* does not retrieve, for example, "compound eye development" even though it has been reported that *lilli* interacts with and regulates *ato* (and *Da*) thus affecting eye development (see main text for references). Therefore, some GO annotations may be incomplete.

**Table 1.**
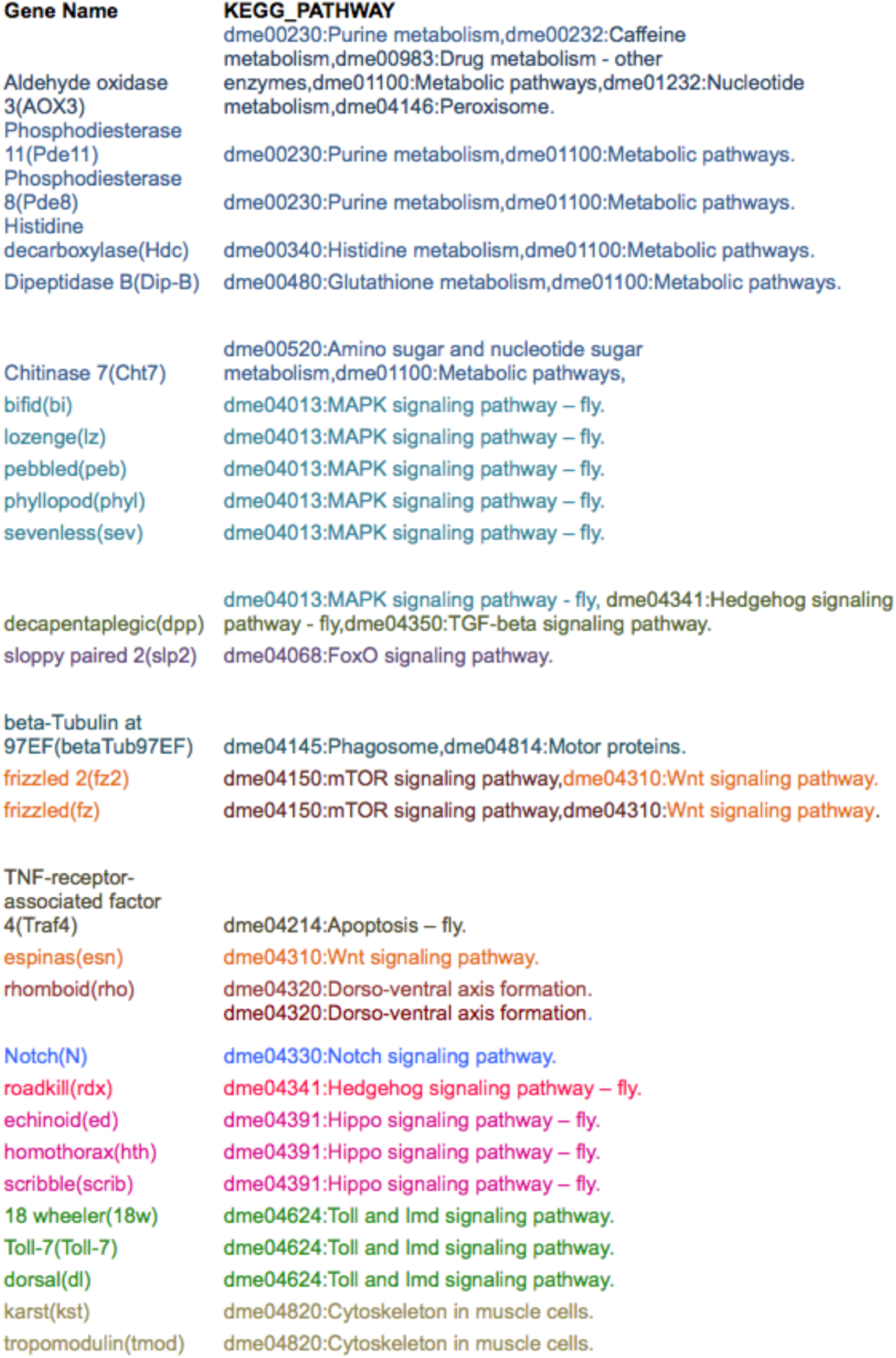
KEGG pathway association of core genes. The gene name (abbreviation) and KEGG pathway code and denomination are indicated.

Of this core set of 106 genes, 22 were TFs (Figure 2A,D,E). These TFs, which we call “kernel” TFs, include all retinal determination gene network genes previously known to be responsible for the early specification and growth of the *Drosophila* eye primordium (*hth, toy, ey, Optix, so, tsh, tio* and *dac –eya* was the only gene of this early acting network not included because it is not a TF, but a transcriptional cofactor of So). *Slp-2* is necessary for the growth of the eye primordium (Sato and Tomlinson, 2007), *gl (glass), peb, lilli* and *luna* are required for the differentiation of retinal cells (De Graeve et al., 2003; Distefano et al., 2012; Moses et al., 1989; Pickup et al., 2002), *B-H1* and *lz* are required for the maintenance of the progenitor cells that feed the retinal differentiation process (Daga et al., 1996; Higashijima et al., 1992), and *oc, lab, Lim1* and *disco* are expressed in prospective head/antennal cells adjacent or surrounding the eye (Diederich et al., 1991; Lee et al., 1991; Roignant et al., 2010; Wieschaus et al., 1992) (Figure 2E). Finally, an involvement of *ovo, snail* or *E5* in the development of the eye disc has not yet been reported. Thus, the identified kernel TF set includes all the known *Drosophila* retinal determination gene network components plus several other genes known to be essential for eye development in *Drosophila*. *Episyrphus* shares these genes, which would be expected to contribute functionally in the same way to eye development in this species. Using the STRING protein functional association database (von Mering et al., 2003) for *Drosophila melanogaster,* the network of the kernel TFs showed high connectivity, confirming a strong functional relationship among them (Figure 2D).

Our results indicated that the *Episyrphus* eye GRN shares many of the known nodes of the *Drosophila* eye GRN. Next, we aimed at finding out how the eye GRN is weaved in both species by predicting regulatory links between TFs and core eye GDARs based on TF binding motifs presence. Of the regulatory kernel of 22 TFs, only 3 lack associated motifs in our collection (those for *bi(omb)*, *dac*, and *tio*). Therefore, the GRN built with the kernel TFs should be very complete.

### DARs of eye genes are enhancers

A gene is recruited as a node in the GRN when an eye-specific DAR is linked to it, since sites of increased chromatin accessibility are good predictors of enhancers (Cusanovich et al., 2018; Janssens et al., 2022; Johansen et al., 2025; Thomas et al., 2011). In any case, we confirmed that this was the case for a small sample of loci encoding TFs in our kernel set in *Drosophila* for which enhancers had been determined by the Flylight project (https://flweb.janelia.org/cgi-bin/flew.cgi; (Jenett et al., 2012)). The *Optix* locus contained four three eye DARs, three of which are eye- and one wing-specific (Figure S4A). Eye-DAR 1 and wing-DAR fell within fragment R30B08, which showed expression in the eye and the wing. Eye-DARs 2,3 fell within fragment R30D11, which showed enhancer activity in two domains in the eye disc: the ocellar region and the compound eye. Likely, each of the two DARs in this fragment is responsible for one of these two activity domains. Therefore, all DARs detected were associated with enhancer activity and in their respective discs. Another example is the intergenic region between paralogs *B-H2* and *B-H1* (Figure S4B). In it, we detected 12 eye DARs. All these DARs, except for three, fell within fragments with eye enhancer activity. Of the remaining three, two fell in regions which were not tested for enhancer activity and the third DAR did so in a region with no imaginal disc enhancer activity reported, but overlapping an annotated long non-coding RNA (Figure S4B). Again, this showed an excellent, almost one-to-one, correspondence between DARs and enhancer activity in *Drosophila.* Together with previous results, this analysis reinforces the idea that DARs generally have an enhancer activity, and activity we extend to *Episyrphus’* DARs. Thereby, these DARs serve as directional anchor points between the regulators (TFs) and the network nodes (GDARs).

### Prediction of TF binding motifs and the eye GRN

Establishing a regulatory link between a TF and a gene depends on the finding of the TF motif in the DARs of the prospective target gene. Therefore, once the collection of eye GDARs had been identified, the next step was the identification of these motifs within DARs. As a method to detect TFBSs in our DARs collection, we used HOMER (http://homer.ucsd.edu/homer/motif/; (Heinz et al., 2010). See Supplementary Methods/GRNs inference/Appendix I for a comparison of motif search methods). In brief, HOMER uses Positional Weight Matrices (PWMs) to represent motifs, with a preset threshold (score_0_) that the target sequence must exceed (Max. score) for the associated TF to be considered as binding. The HOMER motif collection with known score_0_ and directly usable by the search algorithm included only 1005 PWMs (at the time this paper was written), and many of these (636) were discarded because they belonged to TFs without an ortholog in *Drosophila* or to RNA-binding proteins. To broaden HOMER’s motif search capabilities, we integrated various PWM collections, totalling 9860 motifs (Supplementary Methods/GRNs inference/Appendix II). However, all new PWMs lacked a preset score_0_, so these had to be estimated in order to determine the binding criteria for their associated TFs (Supplementary Methods/GRNs inference/Appendix III). Using the collection of PWMs from HOMER with a known score_0_, a predictive model was trained (elastic net linear regression) which, with acceptable predictive capacity (∼60% R²), later enabled the prediction of this threshold for the expanded collection of PWMs. These were the “estimable” set of motifs. However, some of these PWMs had characteristics that differed significantly from those in HOMER, which were used in the model fitting, leading to extreme score_0_ estimates. These PWMs are classified as "not estimable" (3087). However, instead of discarding them, we set their score_0_ to 95% of the theoretical maximum score to make the TF binding predictions more conservative (Figure S5).

With this new collection, we used HOMER to search for motifs of expressed TFs within each eye-specific DARs annotated to nearest gene (GDARs) from both species. Thus, links were established between TFs and target genes. As an overall quality check of the predicted connections, we performed two sets of tests (see Supplementary Methods/GRNs inference/Appendix IV). First, we asked whether motifs for TFs expressed exclusively in the eye were enriched in eye-specific DARs. Indeed, we found this to be the case for some, although not all motifs (Figure S6A). Notably, motifs directly associated with *Drosophila* TFs (that is, not derived from human or mouse orthology) and those deemed "estimable" showed the greatest eye/wing enrichment. Thus, eye/wing enrichment could be used as a measure of confidence in the links established through these motifs. As a second test, we asked if these links complied with the expectation that developmental TFs in the GRN should be, on average, more connected than other genes in the network, as they are often involved in mutual cross-regulation (Marletaz et al., 2018; Sorrells and Johnson, 2015). We used two measures of connectivity. The number of DARs per gene and the number of unique motifs found in all DARs linked to each gene (a sort of upstream regulatory “diversity”). We observed that indeed TFs had more linked DARs as well as a larger number of upstream TF regulators, both in *Drosophila* and in *Episyrphus* (Figure S6B,C).

After these verifications, the eye GRN built with the 106 core genes included 700 shared edges (that is, a motif for the same TF was present in the DAR regions of the orthologous gene in both species), 384 that were *Episyrphus* specific (the motif was present only in the DARs of the *Episyrphus* ortholog) and 390 were *Drosophila* specific. The edges linking a TF to a specific motif in an eye GDAR contained information on: (1) whether the edge had been identified in either of the species or in both; (2) whether the motif derived from *Drosophila* or from human/mouse orthologs; (3) whether the motif was enriched in eye DARs relative to wing DARs, and (4) whether the motif was estimable or non-estimable (see above). With all this information, we generated a GRN model comprising the 106 core eye genes (Supplementary Results/Cytoscape_GRNs_visualization). The GRN for the 22 kernel TFs (19 of which have an associated motif) is presented in Figure S7.

One first observation, both in the *core genes* network and the *kernel TF* network was that the degree of conservation of regulatory links varies between pairs of orthologs. To compute the different degrees of shared connectivity observed, we compared the set of TFs (motifs collapsed by TF) potentially binding the regulatory landscape (sum of associated DARs) of each orthologous gene pair. Using the Jaccard distance as a measure of dissimilarity -that is, two orthologs were deemed regulatory identical (“least dissimilar”) if their DARs had the same collection of binding TFs- we could sort the core genes by degree of shared connectivity (Table S5). On the contrary, they would be most dissimilar if their TF collections were non-overlapping.

Considering just the collection of potential upstream TFs does not reflect the fact that the expression pattern of a gene is usually the result of the piecemeal action of several enhancers, each often active in distinct spatial and temporal subdomains (see, for example, the cases in Figure S4A and B, above). Therefore, comparison of the TF-binding composition of individual enhancers might reveal different degrees of similarity among enhancers, with those most similar being responsible for shared aspects of the gene’s expression pattern, while those most dissimilar conferring species-specific regulation. Thus, we decided to cluster (using hierarchical clustering) the DARs of each pair of the 106 orthologs of the core set, based on their TF-binding composition (using Jaccard distances between pairs). Results are shown as Supplementary Results (*Inter-orthologous DARs clustering panel.pdf*).

### Identification of genes with potentially highly conserved regulation based on DAR distribution, TF motif composition and expression pattern: *glass (gl)* as an example

One example of a gene showing a high degree of predicted regulatory similarity is the TF *gl (glass),* which is expressed and required for the differentiation of the neural and non-neural cell types of the retina (Bernardo-Garcia et al., 2016; Liang et al., 2016; Morrison et al., 2018; Moses and Rubin, 1991). *gl* ranks 100 out of 106 genes in the list of increasing connectivity similarity (Table S5), and it is the most similar among the kernel TFs factors (Table S6). The two orthologous *gl* shared most of their predicted TF regulators (Figure 3A; shared kernel subset). When the two loci were compared, both showed two eye DARs, one overlapping the promoter and another immediately 5’ to it (Figure 3B). Clustering analysis based on predicted TF binding indicated that *Drosophila* eye DAR-1 and *Episyrphus* eye DAR-1,2 were the most similar, sharing binding sites for Optix, Hth, Ovo, So, E5, Limb1, Luna and Lab (Figure 3C,D). Analysis of gene transcription in single cell data shows *gl* coexpression with *so, B-H2, luna, ovo* and *hth* in the differentiating retina population. *Optix* is expressed in eye progenitors and precursors, and is mutually exclusive to *gl,* suggesting a repressive interaction (Figure S8). *E5* and *lab* are expressed in prospective head cells, questioning their relevance as *gl* upstream regulators (Figure S8). These results suggested a very similar regulation of *gl* in *Episyrphus* and *Drosophila.* To test this, we analyzed *gl* expression in *Episyrphus* eye discs using *in situ* hybridization (ISH, Figure 3E). *gl* transcription is detected posterior to the MF, with the highest levels immediately after it, then weakening in more differentiated retinal cells, exactly as has been described for *Drosophila gl* using also ISH (Moses and Rubin, 1991). Therefore, the similarity we detected in *gl* TF regulation and DAR similarity correlated with the high similarity in the expression pattern of the *gl* orthologs.

**Figure 3.**
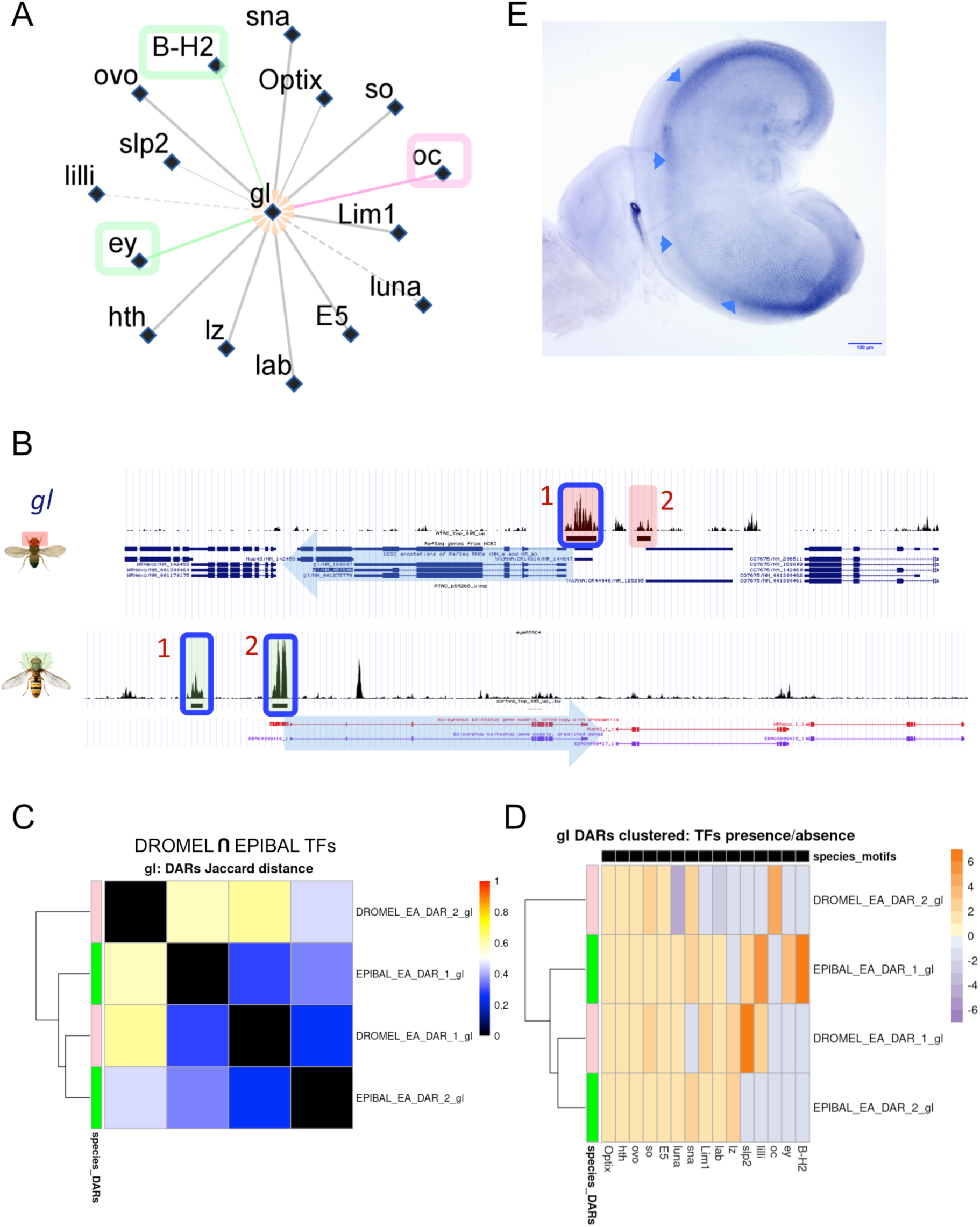
Comparative GRN exemplified by the *glass (gl)* locus. (A) *gl* regulatory network considering the kernel of TFs. Links found in *Drosophila* (“DROMEL”) and *Episyrphus* (“EPIBAL”) are represented in pink and green, respectively. Most links are found in both species (grey), except for Oc/Otd (*Drosophila*-specific) and Ey and B-H2 (*Episyrphus*-specific). Line type: *solid*: DROMEL motifs; *dashed*: human and mouse motifs; Increasing Line *Transparency*: estimable to non-estimable; Increasing Line *Width*: non enriched in eye versus wing to enriched (see Supplementary Methods/GRNs inference/Appendix III for details). (B) Genome browser view of the *gl locus* in *Drosophila* and *Episyrphus*. The transcription unit is marked by the blue arrow pointing to 3’. DARs are shaded and numbered. (C) Clustering of *gl* DARs according to their motif composition similarity (kernel TFs), with colder colors indicating increasing similarity. *Drosophila* and *Episyrphus* DARs are in pink and green, respectively. The most similar DARs are DROMEL DAR-1 and EPIBAL DAR-1,2. (D) Same clustering showing distribution of kernel TF presence. The color warmth is inversely proportional to the frequency of finding the motifs for a given TF in the set of eye-GDARs for each species, and can be interpreted as an indication of how much more atypical a motif is compared to an ubiquitous one. DROMEL DAR-1, and EPIBAL DAR-1,2 all contain motifs for Optix, Hth, Ovo, So, E5, Limb1, Luna and Lab. (E) *gl* expression in *Epysirphus* eye discs, detected by mRNA *in situ* hybridization (ISH). *gl* expression starts posterior to the MF (arrowheads). Immediately after the MF *gl* expression is highest, with the signal becoming weaker more posteriorly. This expression pattern recapitulates that described, also using ISH, in *Drosophila* eye discs (Moses and Rubin, 1991). Scale bar: 100 microns.

### Variation of DAR number, location and TF motif composition between orthologous loci: *dachshund (dac)* as an example

We noted, however, that in general neither the number of eye DARs nor their location in the locus were conserved between orthologous genes. In addition, when similarity in TF motif composition was analyzed using our clustering method, the degree of similarity among DARs linked to orthologous genes varied broadly, although, as a general rule, the degree of DAR similarity was low (See Figure S9 for some examples of orthologs). Among these, only E5 DARs showed a high degree of similarity). Either equivalent regulatory information was distributed differently across *cis*-regulatory regions, or this regulatory information had evolved to be different –although these differences may or may not result in different expression patterns. In the absence of a comprehensive characterization of the expression patterns of the kernel TFs in *Episyrphus,* we began characterizing *dachshund* (*dac*) as an example of divergence in the regulatory predictions of the GRNs. *dac* is a central gene within the *Drosophila* retinal determination gene network: its expression is kept low in eye progenitors but becomes upregulated by the protein complex formed by So and Eya in eye precursors just anterior to the differentiation wave (morphogenetic furrow, MF). *dac* expression remains high straddling the MF and just posterior to it, but then decays as retinal differentiation progresses. In addition, *dac* is expressed in a ring in the adjacent antennal primordium (Bras-Pereira et al., 2015; Mardon et al., 1994). When we examined *dac* expression in *Episyrphus,* using a crossreacting antibody, we detected *dac* expression in an antennal ring as well as in the eye (Figure 4A). In this latter, *dac* was expressed in all cells anterior to the MF, with expression decaying posterior to it (Figure 4A). Therefore, *dac* expression is globally similar in *Drosophila* and *Episyrphus.* Next, we looked among the kernel set of TFs for potential *dac* regulators in our GRN model (Figure 4B). Of these, a subset comprised regulators shared by both species, another subset was *Episyrphus*-specific and finally, *lilli* and *gl* were predicted to be *Drosophila-*specific, although these were low confidence motifs (coming from human/mouse homologs, not eye-enriched and non-estimable). The common predicted regulatory inputs on *dac* were *hth, ey, optix, so,* plus *lz (lozenge), E5, sna (snail), ovo* and *luna* (although this latter is also a low confidence motif). Hth, Ey and So had already been shown to be *dac* regulators (Bras-Pereira et al., 2015; Pappu et al., 2005), and *Optix* was a likely prediction, considering its role in eye development (Li et al., 2013) and its capacity to induce eye development when ectopically expressed (Seimiya and Gehring, 2000). The fact that our GRN model included all known *dac* regulators increased our confidence in other predicted links. Thus, we decided to test one of those novel links: *lz*.

**Figure 4.**
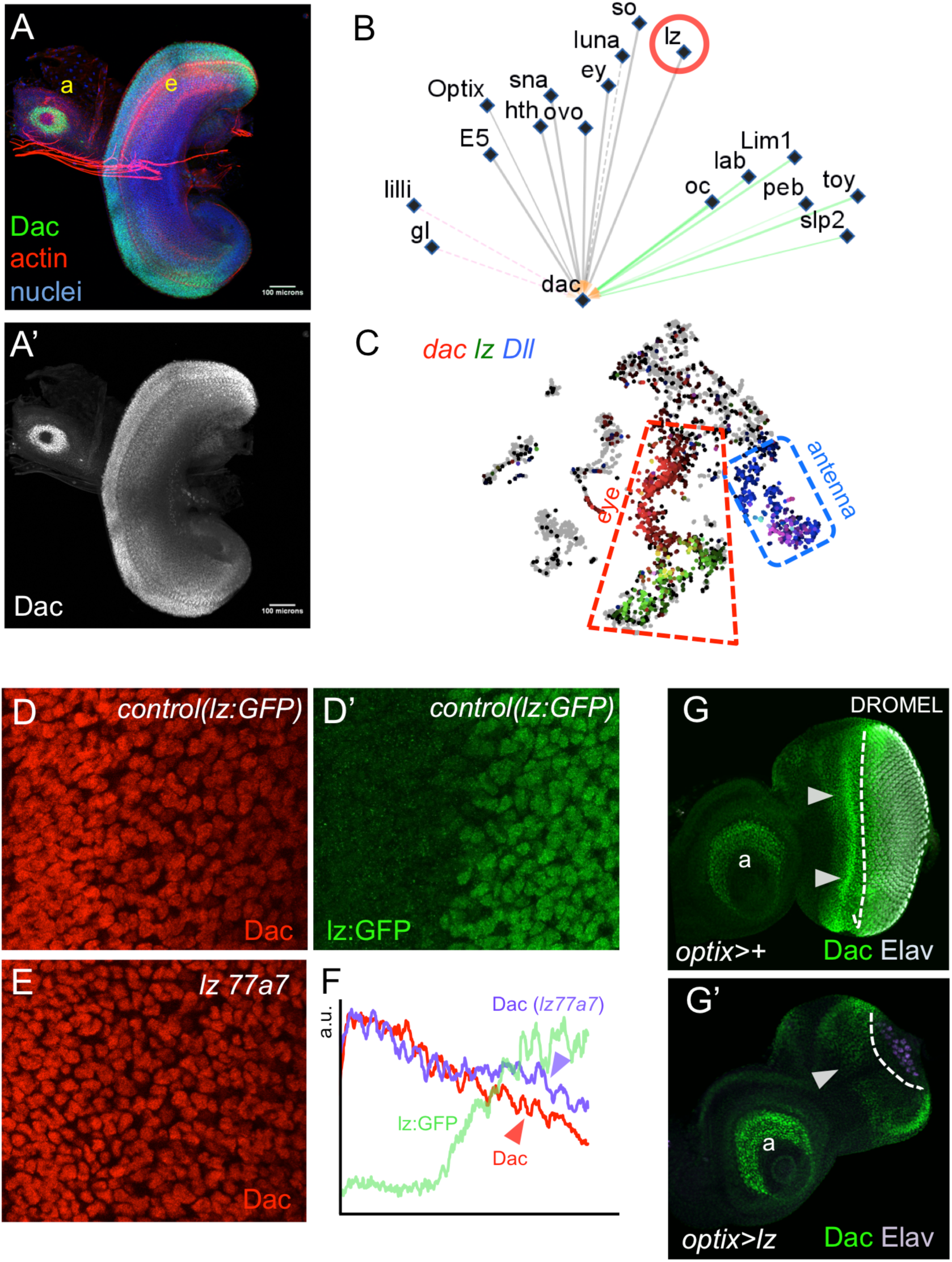
*dac* expression and its upstream regulation. (A,A’) *dac* expression in an L3 *Episyrphus* primordium stained, in addition, for nuclei (DAPI) and actin (phalloidin). The *dac* signal is shown in (A’). the antennal and eye regions are marked by “a” and “e”, respectively. (B) Predicted regulatory inputs on *dac* from the kernel set of TFs. Shared inputs are in grey, *Episyrphus*-specific are in green. *Drosophila-*specific links, which are low confidence, are shown in pink. The TF *lozenge (lz)* is highlighted, as its predicted regulatory action on *dac* is tested in this paper. (C) SCOPE single cell atlas showing the complementary expression of *dac* and *lz.* The eye cells are boxed. The expression of *Dll* is also included to mark the antenna (where *lz* is expressed as well). *Drosophila* eye primordium single cell data is from Bravo Gonzalez-Blas et al., 2020. (D,E) *dac* expression in basal cells in *lz:GFP (Control)* (D) and *lz* mutant *(lz77a7)* in representative *Drosophila* eye primordia. In controls, *dac* expression (Dac, D) decreases as *lz:GFP* expression (*Lz:GFP*, D’) increases. (E) This decrease is less marked in *lz77a7* mutants. (F) Signal quantification of (D, D’ and E). The Dac profile in control and lz77a7 eyes is represented in red and purple, respectively. The *lz:GFP* profile (from D) is shown in green. (G,G’) *dac* expression in control (*optix>+*) and in *lz*-overexpressing eye primordia *(optix>lz)*. The arrows point to *dac* expression in the *optix* domain, anterior to the MF (dashed line). In *optix>lz* (G’) *dac* expression is almost completely repressed, compared to the controls (G). Retinal differentiation, marked with the photoreceptor marker Elav, is also impaired in *optix>lz* primordia. “a”, antenna.

*lz* encodes a TF of the AML1/Runx gene family. In the *Drosophila* eye, *lz* is required posterior to the MF for the differentiation of most retinal cell types, including photoreceptors, lens-secreting and pigment cells (Daga et al., 1996; Flores et al., 1998). Previous work had shown that Dac and Lz expression domains were complementary posterior to the MF (Hayashi and Saigo, 2001), something we confirmed using single-cell transcriptomic data (Figure 4C), suggesting already a *repressive* interaction between *lz* and *dac.* We carried out three experiments to test this regulatory relationship. First, we checked the *in situ* expression of *dac* relative to *lz* detecting Dac in a strain in which *lz* had been GFP-tagged *(lz:GFP).* Lz is expressed in cells whose nuclei lie mostly basal in the epithelium (Flores et al., 1998). Dac expression decays posterior to the MF coinciding with the expression of *lz:GFP* (Figure 4D,D’), confirming previous reports (Hayashi and Saigo, 2001). Second, we analyzed *dac* expression in an eye-null *lz* mutant, *lz77a7* (Flores et al., 1998). In *lz77a7* primordia, Dac expression is maintained at higher levels than in controls in basal nuclei (compare Figure 4D and E), something that is also observed when Dac signal is quantified (Figure 4F), indicating that *lz* is required for *dac* repression. Finally, we ectopically drove Lz anterior to the MF, where *dac* expression is maximal, using the *optix2/3-*GAL4 driver (Ostrin et al., 2006). In *optix2/3-GAL4; UAS-lz* (“*optix>lz”*) eye discs, *dac* expression was dramatically down-regulated (Figure 4G,G’). Therefore, our experiments showed that *lz* is necessary and sufficient for *dac* downregulation thus validating the regulatory link from *lz* to *dac* derived from our GRN analysis, at least, in *Drosophila*.

We next wanted to understand how this regulatory information was distributed in the two fly species. To this, we mapped the identified DARs on the orthologous loci. *Drosophila dac* showed two DARs, one intragenic and another immediately 3’ to the end of the transcription unit, while *Episyrphus dac* had four, all of them 3’ to the end of the transcription unit (Figure 5A). The *Drosophila dac* DAR1 and *dac* DAR2 mapped within the sole *dac* eye enhancers known to date (3EE and 5EE, respectively. (Pappu et al., 2005) and Figure 5), confirming the sensitivity of our ATAC-seq data. When we clustered all six DARs, *Episyrphus* DARs 1, 2 and 4 clustered closer to *Drosophila* DAR1, while *Episyrphus* DAR3 was the most dissimilar among DARs (Figure 5B), although the level of similarity was, in general, weak. In particular, *Episyrphus* DAR1 shared with *Drosophila* DAR1 motifs for *hth, Optix, so* and *lz,* among others (Figure 5C). *Drosophila* DAR1 (3EE; (Pappu et al., 2005) has been shown to be expressed in the developing eye (Pappu et al., 2005) and in the optic lobe, where a gene regulatory network similar to the retinal network is known to operate (Pineiro et al., 2014). We decided to test whether these shared motifs were predictive of eye and optic lobe-specific activity. We cloned the *Episyrphus* DAR1 upstream of GAL4 and examined its enhancer activity in *Drosophila* (*EPIBAL dac_DAR1-GAL4; UAS-GFP*). Indeed, this enhancer showed specific expression in both domains (Figure 5A’). This result indicates that the *Episyrphus* sequence is capable of reading the regulatory state of the *Drosophila* eye and optic lobe correctly, suggesting that a similar regulatory state (that is, a similar set of TFs and active signaling pathways) is present in *Episyrphus.* As we noted above, the motif composition is not particularly similar between *Drosophila dac-DAR1* and *Episyrphus dac DAR1*, and yet, the expression pattern driven by the latter in *Drosophila* recapitulates the expression domains where the former is active. This may indicate that the combination of the shared TFs is sufficiently specific to activate *Episyrphus* DAR1 in the correct territories. Other TF motifs may play secondary or refining roles. Another, non-mutually exclusive explanation, might be that both *Drosophila dac DAR1* and *Episyrphus dac DAR1* shared additional sequences not included in our motif collection, or some sequence “grammar” we have not considered –that is, these DARs would actually be more similar than our method could detect.Globally, these results made three points. First, DARs identified by ATAC-seq in *Episyrphus* also pointed to enhancers, at least when assayed in *Drosophila.* Second, the *trans-cis* code of the regulatory regions (that is, the correspondence between TFs and their target DNA binding motifs) is conserved between *Drosophila* and *Episyrphus,* allowing *Episyrphus’* enhancers to “interpret” the regulatory state of *Drosophila* cells driving a coherent pattern. A recent example of this “coherence” has been reported for *Aedes* enhancers in *Drosophila* transgenics (Schember et al., 2024). And third, the number and synteny (i.e. relative position in the genome) of the *dac* enhancers are not conserved between the two species. As mentioned above, this might be a rather general feature, at least among developmental genes. In the sample analyzed, which comprised in addition to *gl* and *dac,* the TFs *ey*, *Optix*, *bi* (*omb*) and *lz,* as well as the BMP signal *dpp* (Figure S9 and Figure 3,5), the number and location of eye DARs varied between most orthologous pairs. In addition, DARs often differed in TF motif composition, which agrees with the high degree of evolutionary fluidity of CREs previously reported (Buffry et al., 2016; Rebeiz and Tsiantis, 2017). Has this fluidity erased all signs of conservation? Or are there links that have been maintained due to gene network constraints? To answer this question, we compared how orthologous gene pairs cluster according to the similarity in motif composition of their DARs relative to the clustering of orthologs with a random assignation of DARs. In this comparison more orthologous pairs remained clustered than expected by chance (Figure S10 and Supplementary Results/DARs-GDARs_clustering/TFbased), indicating that, at least for *Drosophila* and *Episyrphus,* a certain degree of shared motif composition exists, either deriving from a common ancestral CRE or from functional constraints demanding CREs with specific regulatory activities.

**Figure 5.**
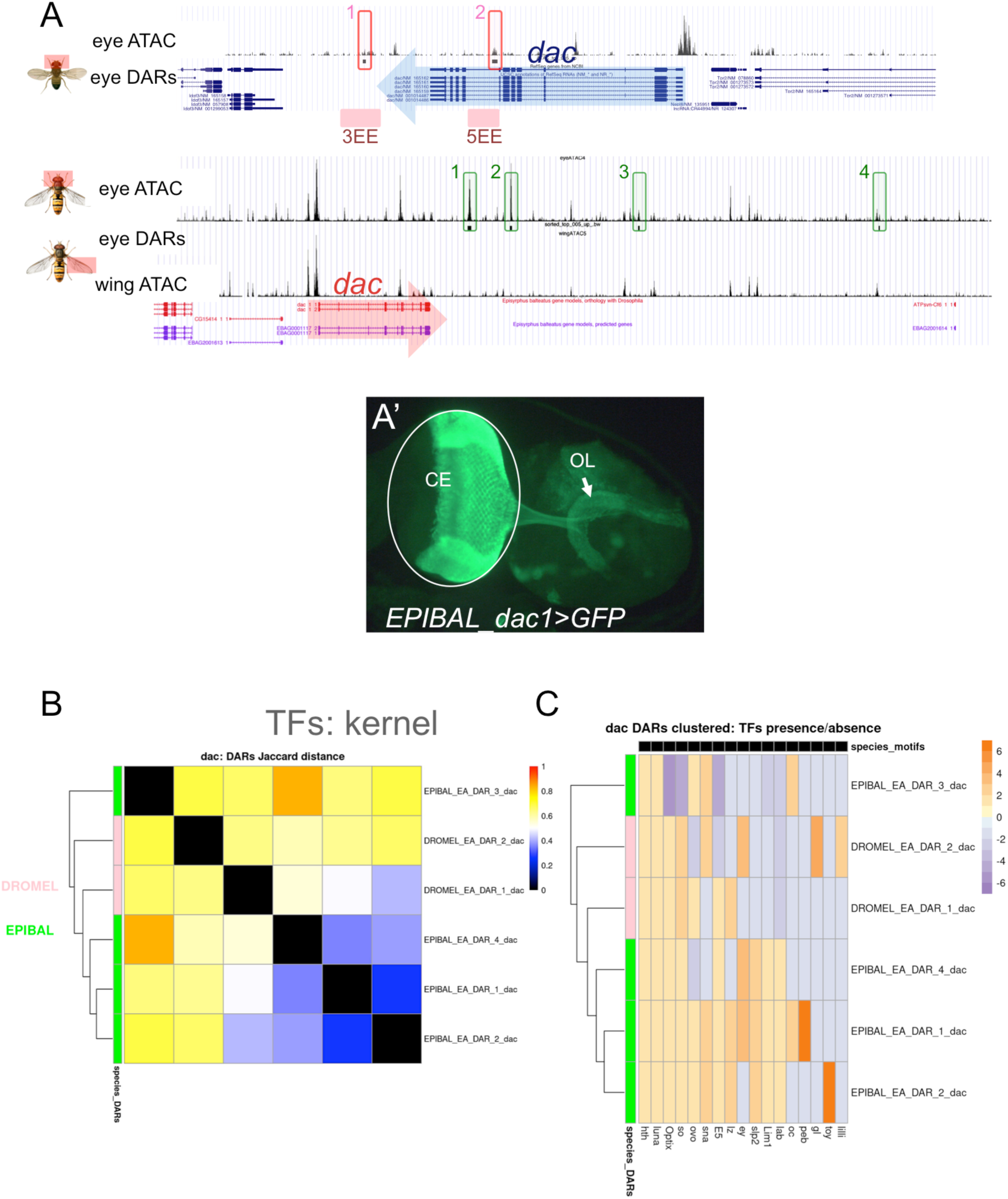
The *dac* regulatory landscape and similarity-based clustering of its potential enhancers. (A) *dac* locus in *Drosophila* (top) and *Episyrphus* (bottom). Genome browser eye tracks (one out of two replicates) are shown. One wing track for *Episyrphus* is also included, to show that these DARs are eye specific. The transcription unit is marked by the blue arrow pointing to 3’. DARs are shaded and numbered. The two *Drosophila* eye DARs correspond to the already characterized 3EE and 5EE enhancers by Pappu et al., 2004 (boxes below the tracks). The ATAC-seq data also identifies clear eye-specific DARs in the *Episyrphus* genome. However, instead of two there are four and, in any case, located in non-syntenic positions relative to *Drosophila,* except for, perhaps, DARs 1 in *Drosophila* and *Episyrphus,* which are located immediately downstream of the transcription unit. (A’) *Episyrphus* dac_DAR1 was cloned upstream of GAL4. Enhancer activity in *Drosophila* transgenes was detected by GAL4-induced GFP expression (see material and methods for details). The expression domains detected are labeled as: CE: compound eye; OL: lamina of the optic lobe (arrow). These are all domains where the endogenous *dac* gene is expressed (Mardon et al., 1994; Pineiro et al., 2014)). (B) Clustering of *dac* DARs according to their motif composition similarity, with colder colors indicating increasing similarity. *Drosophila* and *Episyrphus* DARs are in pink and green, respectively. The collection of motifs used was that for TFs shared by both species (“kernel” set). (C) Same clustering showing the distribution of kernel TF motifs. Color warmth inversely proportional to TF frequency in the whole DAR set (as in Figure 3). Shared kernel TFs between DROMEL dac_DAR1 and EPIBAL dac_DAR1 are: *hth, Optix, so, lz, luna, ovo* and *E5.*

## DISCUSSION

### The comparison of *Episyrphus* and *Drosophila* eye transcriptomes extends the set of eye genes

Most comparative analyses of compound eye development have been carried out either among closely related *Drosophila* species (Buchberger et al., 2021) or between *Drosophila (melanogaster)* and distant species within insects, or even further with chelicerates and myriapods (Baudouin Gonzalez et al., 2025; Baudouin-Gonzalez et al., 2022; Blackburn et al., 2008; Dong and Friedrich, 2010; Prpic, 2005; Samadi et al., 2015; Schomburg et al., 2015; Yang et al., 2009b). In general, long-range evolutionary comparisons involve a small set of genes selected based on prior knowledge derived in *Drosophila.* Here we have aimed at starting to bridge this gap by constructing an eye GRN model in the hoverfly *Episyrphus balteatus* and comparing it to a *Drosophila* eye GRN built using an identical approach.

The first result was the global characterization of genes differentially expressed in the developing eye disc compared to the wing disc. Here, the eye gene sets (“eye DEGs”) point to the same functions in the two species and match perfectly the developing eye processes (Figure 1). However, even though both eye gene sets are characterized by similar functional and molecular terms, there is only limited overlap between them (158 shared eye DEGs). That exclusive *Episyrphus* eye genes are annotated as “eye-related” comes from the fact that these same genes (orthologs) have been annotated as such in *Drosophila*. Then, why weren’t these *Episyrphus* exclusive “eye-related” genes identified as eye genes in *Drosophila* too? One explanation is that genes annotated with eye development-related functions might as well be expressed in the wing disc, so that some of the “eye genes” were left out after the eye versus wing comparison. If the wing discs of the two species differed in the expression levels of some eye genes, the resulting eye DE gene sets would also differ. Since the process of photoreceptor differentiation is exclusive to the eye, we expected this 158 gene set to be particularly enriched in genes expressed specifically in retinal cells, which was indeed the case: 34% of them were associated to Nervous System Development (Figure 1G,H)–which, in the eye-antennal primordium mostly represents retina development and differentiation. Therefore, the set of 158 shared eye DEGs (Figure 1) should be considered a conservative list of genes specifically associated to the development of the compound eye across a broad swath of the Diptera tree of life.

### Some considerations on our GRN-building strategy

A critical step for GRN modeling was the identification of eye-specific CREs and their linked genes, as these CREs would be the primary sites of GRN control. As we described above, we identified eye DARs (that is, ATAC peaks that were detected in eye but not in wing discs) and linked them to the closest eye-expressed gene. This approach has a number of caveats. A first caveat is that of the limits of a regulatory locus. While the association between an intragenic DAR and the hosting locus is usually safe, the association of an intergenic DAR to any gene upstream or downstream to it may depend on where the chromatin boundaries lie (Oudelaar and Higgs, 2021), so that in addition to proximity, the association should take into account boundaries, something we did not do, as this type of data was not available for *Episyrphus.* However, GO analysis of the genes linked to eye-specific DARs indicated that, globally, the associations should be correct, since the enriched terms clearly pointed to eye development in both *Drosophila* and, especially, in *Episyrphus.* In addition, we found enrichment of motifs for TFs expressed in the eye in eyeDARs, when compared to wing DARs, especially for motifs identified for *Drosophila* TFs, globally supporting the connectivity of our network model.

The second caveat affects the sensitivity of DAR detection. Although the probability of missing a DAR in a bulk experiment increases with the rarity of the cell state within the tissue in which the enhancer is active, we were aiming at the gene network responsible for the specification and growth of the eye and the initiation of its differentiation. During these processes, the relative proportion of cells in the same state is expected to be large and therefore, it is also very likely that many cells are activating the same CREs (that is, opening their regulated DARs), making them readily detectable using ATAC-seq. This is why we expected that, by and large, we should not be missing major enhancers in neither of the two species. The *Drosophila* gene sample for which we studied the correspondence between functional enhancers and DARs, although very limited, showed almost biunivocal correspondence, supporting the idea that our ATAC-seq data was not missing major enhancers in *Drosophila,* which led us to assume that likely neither it did in *Episyrphus*.

### A conserved eye GRN integrated in the global regulatory network of the developing head

With all these considerations in mind, we built the GRN model with the 106 eye GDARs (cytoscape file available in Supplementary results/ Cytoscape_GRNs_visualization). In Figure 6 we summarized a large part of this network. In it, we show the eye GRN comprising the core genes expressed in the developing eye (as in the single cell atlas by (Bollepogu Raja et al., 2023)). These genes (nodes) were located more or less centrally depending on their degree of connectivity, with their size representing their degree of regulatory similarity (with larger nodes being more similar). In addition, we include temporal information on their expression based on the single cell atlas of *Drosophila* eye disc produced by the Mardon laboratory (Bollepogu Raja et al., 2023). We further assumed, as a first approximation, that the same temporal dynamics applied to the *Episyrphus* orthologs.

**Figure 6.**
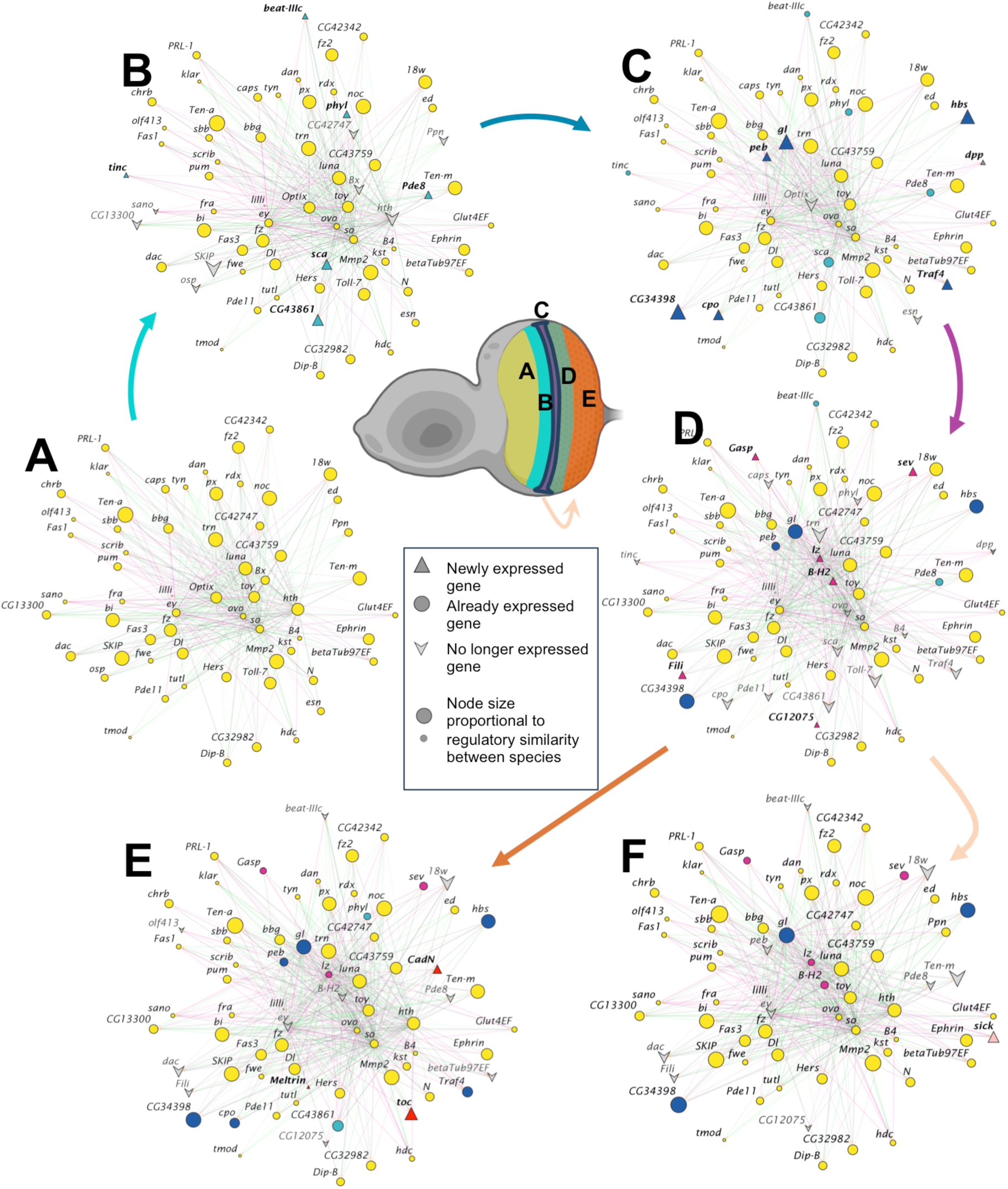
The collapsed eye GRN containing the core gene set and its predicted regulatory interactions. The network represents the changes in gene expression as they occur during *Drosophila* eye development (see Materials and Methods/*Identification of core genes expressed in each eye primordium cell population using single-cell RNA-seq*). (A) Eye progenitor; (B) Precursor; (C) morphogenetic furrow region; (D) early photoreceptor; (E) late photoreceptors and accessory cell types; (F) basal retinal progenitors. Colour and line type of links as in Figure 3.

When kernel TFs were considered, the resulting network was highly interconnected, in agreement with the retinal GRN being subject to many regulatory feedbacks. Of all kernel TFs, only *sna* and *ovo* lack evidence of being involved in eye disc development. Our network includes all the known *Drosophila* retinal determination gene network (RDGN) (*ey, toy, so/Six2, Optix/Six3* and *dac*) and extends these cross-regulations to include further TFs, involved in stimulating the growth of eye progenitors (*hth, tio* and *tsh*) or in driving retinal differentiation (*gl, peb, lilli, luna, disco, lz, B-H1, B-H2*). Some of the kernel TFs have been previously involved in the development of disc derivatives other than eye, like the head (*oc, lab, Lim1, slp2*). Since the starting material was the eye-antennal disc, this was expected. The fact that we predicted them connected within the network might indicate regulatory relationships between the genetic programs controlling the eye and other regions of the disc, like the mutual repression between *ey* and *cut,* which has been proposed previously to be responsible for the partitioning of the disc into eye and antennal fields, respectively (Ramaekers et al., 2019; Wang and Sun, 2012). Therefore, the eye GRN should be considered a part of a global gene regulatory network controlling the development of all the head structures from *Episyrphus* to *Drosophila* at least. In addition, we identified four cases, in which the kernel TF (*tio, B-H2, E5* and *slp-2*) was part of a linked paralogous gene couple: *tio/tsh, B-H2/B-H1, E5/ems,* and *slp-2/slp-1.* It has been previously shown for these pairs that paralogs had very similar, if not identical expression patterns (Bessa et al., 2009; Datta et al., 2009; Lim and Choi, 2003; Mahaffey et al., 2001; Sato and Tomlinson, 2007). In the case of *E5* and *ems* single cell RNAseq data showed also coincidental expression (Figure S3A,B). Therefore, the corresponding paralogs may well be part of the conserved eye GRN.

### Evolution of CREs: conservation and divergence in orthologous loci of developmental genes

The comparative analysis of CREs between *Drosophila* and *Episyrphus* reveals significant variability in the conservation and organisation in the orthologous loci of developmental genes. While some genes, such as *gl*, exhibit highly shared connectivity, others, such as *dac*, show greater divergence, suggesting varying degrees of evolutionary flexibility in gene regulation. This variability in shared connectivity implies that not all genes in the GRN are equally free to evolve, which could influence the stability or plasticity of their expression patterns. However, we did not identify any rule that could help us predict the shared connectivity observed. This metric (shared connectivity) showed no correlation with the degree of connectivity and remained invariant across gene ontologies (not shown).

A particular point of interest is the *Drosophila* enhancer *dac* 3EE and its potential counterpart in *Episyrphus* (DAR-1). Despite the low similarity in motif composition, DAR-1 exhibited regulatory activity in the eye and optic lobe lamina of *Drosophila*, similar to that of 3EE, suggesting that only a subset of the TFs predicted as regulators might be truly functional, with the rest being redundant or subtle modulators of expression. However, since this assay was conducted in *Drosophila*, the result depends on the TFs expressed in the developing eye of this species. To confirm that DAR-1 in *Episyrphus* exhibits equivalent activity, functional assays in its native context would be required.

More broadly, the comparison of orthologous loci suggests that the distribution of CREs has shifted over the 90 million years of evolution separating the two species. The lack of conservation in sequence and arrangement of the DARs supports the idea that regulatory information is highly dynamic and can be rearranged without necessarily altering the expression patterns they control. This reinforces the notion that regulatory evolution is not a rigid process but rather fluid, generating diversity without compromising the functionality of developmental programs.

### The extended, conserved GRN model allows making testable predictions about gene regulation

The final GRN model included a set of shared 106 eye genes connected by conserved and species-specific regulatory links. Our criteria for calling the presence of a TF motif were relaxed (for example, we did not include any motif “grammar” as criterion). However, we observed an enrichment in TF motifs for eye-specific TFs especially when the motifs were derived from *Drosophila,* supporting the specificity of the predictions. An additional criterion strengthening the confidence on predicted links could be conservation, especially since we found that, as a general rule, motif composition was not particularly conserved when DARs from paralogous genes were compared. We tested this hypothesis for a new predicted regulatory link between *dac* and *lz.* Our results, including loss and gain-of-function experiments, showed that indeed *lz* was a *dac* regulator in *Drosophila*, confirming the predicted regulatory relationship. The fact that Lz binding sites were found on *dac* DARs suggested that this regulation is direct. We think it likely that this regulatory link also exists in *Episyrphus.* But how generalizable are these GRN predictions? Testing this sort of hypotheses (that is, whether predicted regulatory links and indeed functional) will require further genetic tool development in *Episyrphus* -as these should allow manipulating gene function post-embryonically. Even in *Drosophila,* a systematic testing of these hypothetical links, being these tests biochemical (e.g. chromatin immunoprecipitation) or genetic (e.g. mutational analysis, RNA interference), would not be straightforward. In other words, a systematic functional analysis of a GRN might not ever be practical and always remain a prediction, at least partially. It is likely that deep learning methods using large collections of data may be used to increase the probability of predicting true positive links. However, we foresee that GRN models will most likely be tested not at the level of individual links, but by comparing the global effects of specific perturbations *in vivo* and *in silico*, such that good agreement between biological and computational predictions would validate the GRN model as a whole.

## MATERIALS AND METHODS

### Episyrphus balteatus culture

*Episyrphus* larvae and adults are reared in different rearing cages. To feed the larvae, aphids are cultured on broad bean plants in an independent cage. All these cultures were maintained at a temperature of 22±1 °C.

Adults are fed ad libitum with two types of food simultaneously: pollen crushed in a kitchen grinder, spread on paper; and cotton wool soaked in honey dissolved in water. Then there is the egg and larval cage. Larvae feed mainly on aphid-infested bean plants. The broad beans are sown every 2 days, and the aphids from the older plants are then transferred to the newly sprouted plants, which are most easily colonized by aphids. For egg-laying, we place one of these plants in the adults’ cage. After a short time, females will start laying eggs in the places where aphids are concentrated. After a variable time, the plant is removed from the adult cage and placed in a separate one, which is the specific larval cage. This is the time 0 used to measure age, in hours after egg-laying (hAEL). At 36-48 hPEL, the eggs hatch and the larvae start feeding on the aphids. Then, every 48 hours, the larvae are transferred to new, aphid-infested plants. After about 260 hours (10-11 days), the larvae have reached the end of the larval stage and form pupae, usually outside the feeding area. The pupae are collected and placed in a petri dish, which in turn is placed in the adult cage.

### Genomes

*Drosophila melanogaster* genome version dm6 (Hoskins et al., 2015). *Episyrphus balteatus* genome and annotation (Doyle et al., 2022).

### RNA Sequencing and Bioinformatic Processing

#### Experimental Design

RNA sequencing (RNA-seq) experiments were conducted on *Drosophila melanogaster* (DROMEL) and *Episyrphus balteatus* (EPIBAL) to investigate gene expression differences in eye and wing imaginal discs. EPIBAL cultures were raised as indicated above and DROMEL at 25 °C under standard conditions. To stage the discs such that tissues were at an equivalent developmental stage in both species, we used the morphogenetic furrow (MF) position as a marker of differentiation progression. A stage in which the MF had progressed midway along the anterior-posterior axis was chosen, as in these discs an equivalent representation of progenitor/precursor cells (anterior to the MF) and differentiating retina (posterior to the MF) was included for both species. This developmental stage is reached at 96 hours after laying (hAEL) in DROMEL and at 120 hAEL in EPIBAL in our culture conditions.

In DROMEL, samples were collected at two developmental time points: 96 and 114 hAEL, with two and three biological replicates per tissue, respectively. In EPIBAL, two biological replicates per tissue were collected at 120 hAEL. RNA sequencing was performed using the Illumina NovaSeq 6000 platform with a read length of 51 base pairs (bp) in a single-end mode. Libraries were prepared using the Illumina TruSeq RNA Sample Preparation Kit.

#### Sample Collection and RNA Extraction

For each condition, 50 eye or wing imaginal discs were dissected from *Drosophila melanogaster* and *Episyrphus balteatus* larvae at the specified developmental time points. Total RNA was extracted using the RNeasy Kit (Qiagen) following the manufacturer’s instructions. RNA quality and concentration were assessed using a Bioanalyzer (Agilent Technologies) before proceeding with library preparation.

#### Library Preparation and Sequencing

RNA libraries were prepared according to the Illumina TruSeq RNA protocol. The resulting libraries were sequenced on the Illumina NovaSeq 6000 system in a single-end mode with a read length of 51 bp. The sequencing output was quality-checked using FastQC.

#### Bioinformatic Processing

Raw sequencing reads were processed for quality control and filtering using Trimmomatic and Fastp to remove low-quality bases and adapter sequences. Filtered reads were aligned to the reference genome using HISAT2. The resulting alignments were used to generate read count matrices with HTSeq2. Differential gene expression analysis was conducted using DESeq2 to identify genes with significant expression differences between conditions.

#### Additional RNA-Seq Data Acquisition

To complement our experimental data, additional RNA-seq datasets were retrieved from the Gene Expression Omnibus (GEO) repository under accession number GSE59707. These datasets included RNA-seq samples from eye-antennal and wing imaginal discs of wild-type *Drosophila melanogaster* strains (RAL-208 and Canton-S). Raw sequencing data were accessed from the Sequence Read Archive (SRA) under the study identifier SRP044761.

#### Processing of External Datasets

The external RNA-seq datasets underwent the same bioinformatic pipeline as our experimental samples. Reads were filtered using Trimmomatic and Fastp, aligned to the dm6 genome with HISAT2, and read counts were generated using HTSeq2. Differential expression analysis was performed using DESeq2 to ensure consistency with our experimental analyses.

### ATAC Sequencing and Bioinformatic Processing

#### Experimental Design

ATAC sequencing (ATAC-seq) experiments were conducted on *Episyrphus balteatus* (EPIBAL) to investigate chromatin accessibility differences in eye and wing imaginal discs. EPIBAL cultures were raised at 25 °C under standard conditions. To ensure an equivalent developmental stage, we used the morphogenetic furrow (MF) position as a marker of differentiation progress. Larvae were dissected when the MF had divided the disc into two halves, indicating that 50% of progenitors had already differentiated into photoreceptors. This developmental stage is reached at 120 hours after laying in EPIBAL.

ATAC-seq samples were collected at 120 hours after egg laying, with two biological replicates per tissue. Sequencing was performed using the Illumina NovaSeq 6000 platform in paired-end mode with read lengths of 49/25 bp. Libraries were prepared using the Nextera DNA Library Prep Kit.

#### Sample Collection and Nucleic Acid Extraction

For each condition, 7 eye or 21 wing imaginal discs were dissected from *Episyrphus balteatus* larvae at the specified developmental time point. Total chromatin was extracted following the standard ATAC-seq protocol.

#### Library Preparation and Sequencing

ATAC-seq libraries were prepared using the Nextera DNA Library Prep Kit. The resulting libraries were sequenced on the Illumina NovaSeq 6000 system in paired-end mode with read lengths of 49/25 bp. The sequencing output was quality-checked using FastQC.

#### Bioinformatic Processing

Raw sequencing reads were processed for quality control and filtering using Trimmomatic and Fastp to remove low-quality bases and adapter sequences. Filtered reads were aligned to the reference genome using HISAT2. Peak calling was performed using MACS2 (Model-based Analysis of ChIP-Seq). Each tissue conservative peaks were selected using the Irreproducible Discovery Rate (IDR) method, with a global IDR threshold of ≥1. Then, the tissue conservative peaks were merged. Read count matrices for chromatin accessibility were obtained using HTSeq2, and differential accessibility analysis was conducted using DESeq2.

#### Additional ATAC-Seq Data Acquisition

Additionally, DROMEL ATAC-seq experimental data were retrieved from the GEO repository (GSE101827 series). *Drosophila melanogaster* cultures and sampling conditions were consistent and compatible with our omics experiments. The dataset included samples GSM2716860–GSM2716867. Raw data were retrieved from the SRA repository (SRP113497).

#### Processing of External Datasets

The external RNA-seq datasets underwent a very similar bioinformatic pipeline as our experimental samples. Raw sequencing reads were processed for quality control and filtering using Trimmomatic and Fastp to remove low-quality bases and adapter sequences. Filtered reads were aligned to the dm6 genome using HISAT2. Peak calling was performed using MACS2 (Model-based Analysis of ChIP-Seq). Each tissue sample was merged and split into two pseudoreplicates. Subsequently, conservative peaks were selected using the Irreproducible Discovery Rate (IDR) method, applying a global IDR threshold of ≥1. Then, the tissue conservative peaks were merged. Read count matrices for chromatin accessibility were obtained using HTSeq2, and differential accessibility analysis was conducted using DESeq2.

#### DARs annotation

We used the *annotatePeaks.pl* script from the HOMER software [REF] to annotate DARs to the nearest Transcription Start Site (TSS) within a 1 Mb range. The annotations were then filtered to retain only genes with expression levels above a threshold, determined by the median FPKM values across species and tissue samples (see Supplementary Methods/Appendix VI).

#### Motif search

Different motif search methods were discussed (Supplementary Methods/Appendix I) before agreeing to adapt the HOMER method (Heinz et al., 2010) to an expanded collection of motifs (PWMs) derived from three general databases: HOMER, JASPAR24 (Rauluseviciute et al., 2024), and SCENIC+ (Bravo Gonzalez-Blas et al., 2023) (Supplementary Methods/Appendix II). The adapted method involves predicting the binding prediction thresholds used by HOMER (score_0_) for the expanded collection (Supplementary Methods/Appendix III). The predicted score_0_ for each PWM was used as a motif detection threshold on the set of DARs from the eye and wing in both EPIBAL and DROMEL. To this end, the "annotatePeaks.pl" function of HOMER was executed. Finally, the HOMER motif search method applied to the expanded collection was evaluated based on the results obtained from the eye and wing DARs in both species (Supplementary Methods/Appendix IV), revealing differential connectivity of TFs in the eye networks of both species and an eye/wing enrichment of motifs corresponding to genes expressed in the eye of each species.

#### DARs clustering

Both the DARs and the GDARs, obtained by merging the former for each gene, were classified based on the Jaccard distance of their motif composition, following a hierarchical clustering algorithm (Supplementary Methods/Appendix V). The classification results for each subset of motifs used are included in the Supplementary Results, as indicated in Supplementary Methods/Appendix V.

### DAR-GAL4 construct generation and transgenesis

Genomic DNA extraction was carried out according to the protocol obtained from the Vienna Drosophila Resource Center (VDRC) at Vienna BioCenter Core Facilities (VBCF). For *E. balteatus* DNA extraction, three random female adults were picked, and after wings and abdomen removal the rest of the protocol was followed as-is. The protocol can be found at https://shop.vbc.ac.at/media/pdfs/GoodQualityGenomicDNA.pdf.

For cloning, we performed PCR on the gDNA synthesized using the Expand High Fidelity PCR System (Roche) and the amplified band cloned into the pBPGUw plasmid (Adgene plasmid #17575) by Gateway Cloning, following the provided protocols as-is. These DAR containing plasmid were chemically transformed into competent Match1 *E. coli* strain by standard methods and verified by Sanger sequencing. Once verified, transgenes were generated by site-directed transgenesis by FlyOrf (Zurich, Switzerland).

Primers used for the amplification of *Episyrphus dac_DAR1:*

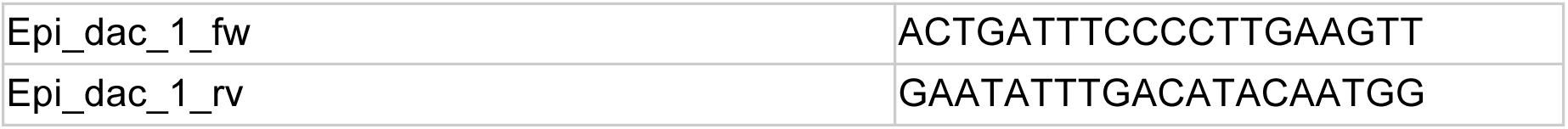

### Cytoscape network models

To compare the EPIBAL and DROMEL RDGRNs, we used the network visualization tool Cytoscape (Shannon et al., 2003). Cytoscape generates a regulatory network by mapping all connections between the source (TFs) and the targets (any gene regulated through its CREs/DARs), while also allowing us to highlight the properties of these connections or the nodes themselves.

For a more comprehensive analysis, the species-specific networks were then superimposed, creating a new network that distinguishes between common and species-specific nodes and connections. Additional information determining the confidence of each motif supporting a connection (Supplementary Methods/Appendix IV) was also incorporated, including the origin of the motifs (DROMEL or Human/Mouse), predictability (estimable or inestimable), and tissue enrichment (eye-enriched or non-significant).

Furthermore, to synthesize the biological and regulatory information, all motifs associated with the same TF were condensed into a single representation. To retain the relevant confidence information for these connections, we adopted a criterion that always considers the ‘best’ properties of the set of motifs associated to the same TF in each species: DROMEL, estimable, and eye-enriched.

These visualizations allow us to observe the RDGRNs of both species on a broad scale or, alternatively, to explore the differential connectivity of specific genes or gene groups, providing valuable insights for generating hypotheses about eye regulation in dipterans.

### Drosophila genetics

The following stocks were used: *UAS-lz* (FBst0033836)*, lz:GFP* (FBst0043954)*lz77a7* (FBst0003609) are described in Flybase (https://flybase.org/) and *optix2/3-GAL4* in (Neto et al., 2017; Ostrin et al., 2006). Larvae from the *optix2/3-GAL4* females to *UAS-lz* males, raised at 25°C, were used to analize the effects of *lz* overexpression on *dac* expression.

### Immunostaining and imaging of *Drosophila* discs

For immunostaining and confocal microscopy, *Drosophila* eye discs were dissected from late third instar larvae in cold PBS and processed essentially as in (Casares and Mann, 2000). Primary antibodies used were: mouse anti-Dac 1-1 (1/50) and rat anti-Elav 7EBA10 (1/1000), both from the Developmental Studies Hybridoma Bank developed under the auspices of the NICHD and maintained by The University of Iowa); rabbit anti-GFP (A11122, Molecular Probes) 1/1000. Appropriate fluorescent Alexa-conjugated antibodies were used at a dilution of 1/500 to detect the primary antibodies. Imaging was carried out on a Leica SPE confocal setup. Images were processed with FiJi (Schindelin et al., 2012).

### *In situ* hybridization for *E. balteatus* glass

#### Identification of the *Episyrphus balteatus glass (gl)* and probe generation

The first step was to identify the orthologous E. balteatus *glass (gl)* gene. To do this, we ran a tblastn of the *D. melanogaster gl* amino acid sequence (Uniprot: P13360 - GLAS_DROME) against the *E. balteatus* genome. The result gave the highest overall score for a gene annotated as *gl_2_1*. We extracted the predicted CDS sequences of this gene, spliced them and translated them into different open reading frames (ORF) using the Expasy Translate tool. We took the ORF which was the size of all the CDSs combined together and applied Motif Scan motif prediction to this amino acid sequence. The predicted protein was very similar to *D. melanogaster gl*, with a series of C2H2 zinc finger motifs at its C-terminal end. This suggests that *gl_2_1* is the *E. balteatus* ortholog of *D. melanogaster gl*.

Next, we generated the *gl* sense and antisense probes (gl_S and gl_AS, respectively) by double PCR. For this purpose, we designed candidate primers flanking several CDS of the gene, specifically exons 3-5. These exons are:

Forward (fw): <u>GGCCGCGGCGG</u>CTGTCTATGTTGCCAGGGGA
Reverse (rv): <u>GCCCCGGCC</u>GGTTGCATGTTGAAGGAGCA

The resulting PCR product was of ∼400 bp. The underlined regions correspond to a universal sequence that was used to generate a cDNA containing our probe with the T7 transcriptase binding site. The non-underlined regions are those that would align during the PCR: in a first PCR, they amplified the *gl* cDNAs of retrotranscribed cDNA from larval plus adult E. balteatus tissues. In a second PCR, we combined our glass primers with universal primers to generate the sequences to be transcribed into gl_S and gl_AS. These 5’ and 3’ universal primers contain the universal sequence complementary to our gl fw and rv primers, respectively, plus a T7 RNA polymerase binding site. Finally, the product of the second PCR was transcribed using a New England biolabs T7 kit (M0251S) and DIG-labelled nucleotides, then precipitated, purified and dissolved in double-distilled water with the addition of 0.5 microlitres of RNAase inhibitor. The quality and integrity of the probes were confirmed by running them on a 2% agarose gel at 120V for 15 min, observing a single band of appropriate size and without smears, and measuring the concentrations in a Nanodrop. The products showed an absorbance curve suitable for a purified RNA. Finally, the probes were diluted in 1 ml final volume of hybridization buffer (sol. A) and stored at −20°C.

### In situ protocol

The *in situ* hybridization protocol used is a variant of the standard alkaline phosphatase (AP) protocol for *Drosophila* imaginal discs (Nagaso et al., 2001). Unless otherwise stated, all washes were 5 min at RT. The major key modifications were the dissection procedure of *Episyrphus* larvae and during the Pre-hybridization step, which we describe in what follows:

#### *Episyrphus* dissection

Third larval stage larvae are collected in wells containing 1× PBS on ice. After a few minutes, when completely still, they are dissected with dissecting forceps, cut just anterior to the level where the posterior fat body is seen under the cuticle. Then, this anterior “cone” is grasped by the mandibles and turned inside-out to expose the imaginal discs and other internal tissues. The posterior part of the larva may or may not be discarded, depending on whether the genital disc is required for sexing each individual. Alternatively, the larva can be partially opened, leaving the anterior and posterior parts together, and both turned inside out. In this way, the genital disc and the rest of the imaginal discs of the same individual can be kept together when sexing is required, and several larvae can be worked with in the same well. After turning inside-out, the digestive tube is removed and the larvae are fixed in 4% formaldehyde in PBS for 25 min at room temperature (RT).

#### Pre-hybridization

After fixation, the samples are washed in PTwx (1x PBS with Triton X-100 and 0.1% Tween 20 each) and twice in ethanol. They are then incubated in 1:1 v/v xylene/ethanol solution for 1 h at RT. This is followed by 2 washes in 100% ethanol. Then, the tissues are washed in serial dilutions of 80, 50 and 25% methanol/ddH20 and finally washed 2 times in ddH20. At this point, the discs are incubated in 80% acetone in ddH20 at −20°C for variable times depending on the age of the larvae (measured as hours (h) after egg laying, hAEL): a) <170hAEL = 10 min, b) 170-200hAEL = 15 min, c) 200-250hAEL = 20 min. After this incubation, the samples are washed 2 times in PTwx, postfixed in 4% formaldehyde for 25 min at RT and washed again 2 times in PTwx.

## ACKNOWLEDGEMENTS

We thank M. Irimia (GRG/ICREA Barcelona) for help with the annotation and orthology predictions of the *Episyrphus balteatus* genome version used in this paper. This work was funded by grants PID2021-122671NB-I00 (AEI, Spain) to FC. TN was a recipient of doctoral contract BES-2016-079458 (AEI; Spain).

## Supplementary Figures

**Figure S1.**
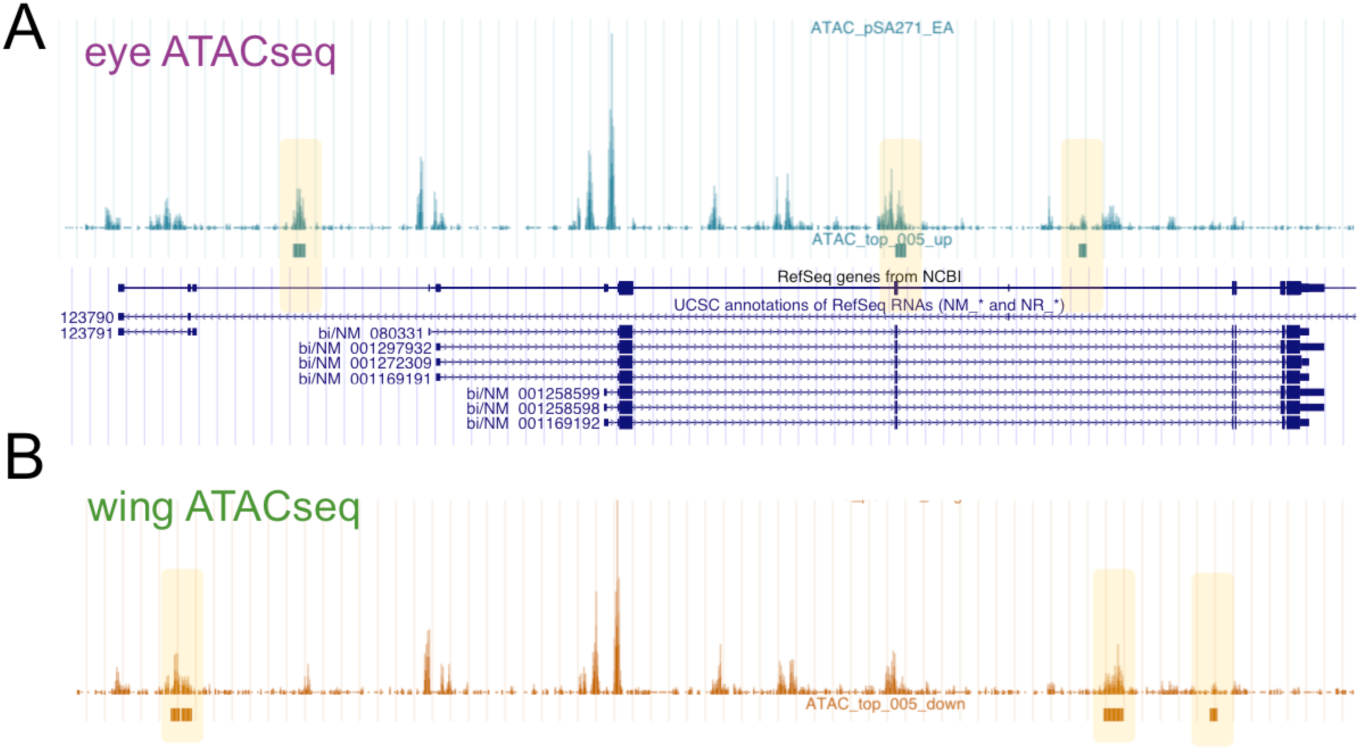
Browser view of the *Drosophila bi/omb* locus. Eye (A) and wing (B) ATACseq tracks are shown above and below it, respectively. Eye-wing differential DARs are indicated as boxes under their respective tracks. While *bi/omb* is not detected as differentially expressed between eye and wing discs, it is linked to eye- and wing-specific DARs indicating this gene is subject to organ-specific regulation.

**Figure S2 (Suppl. To Figure 2).**
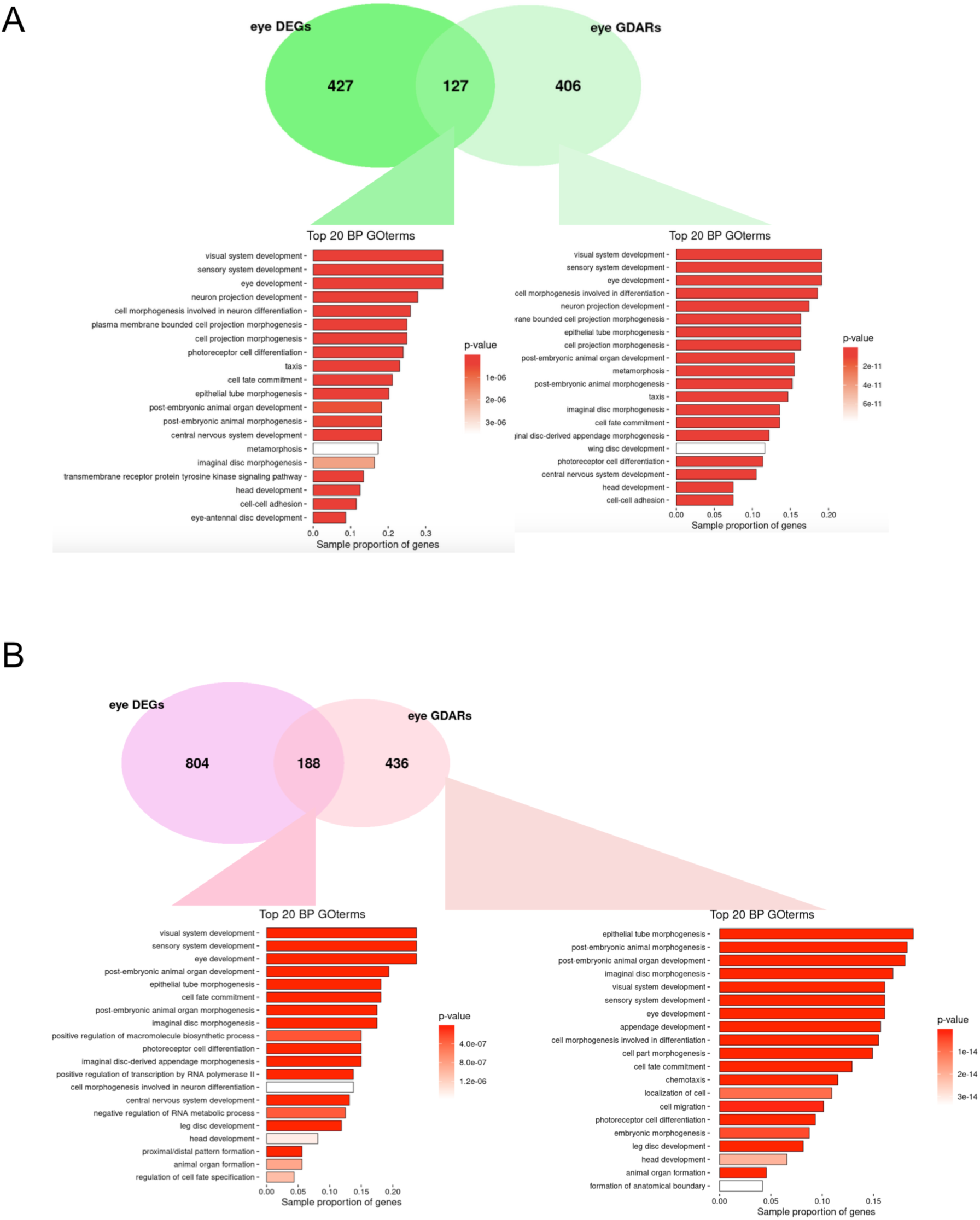
GO analysis (Biological Process, BP) of the eye GDARs in Episyrphus (A) and Drosophila (B) and of the intersection with the DEG set. Top 20 BP terms are shown.

**Figure S3 (Suppl. to Figure 2).**
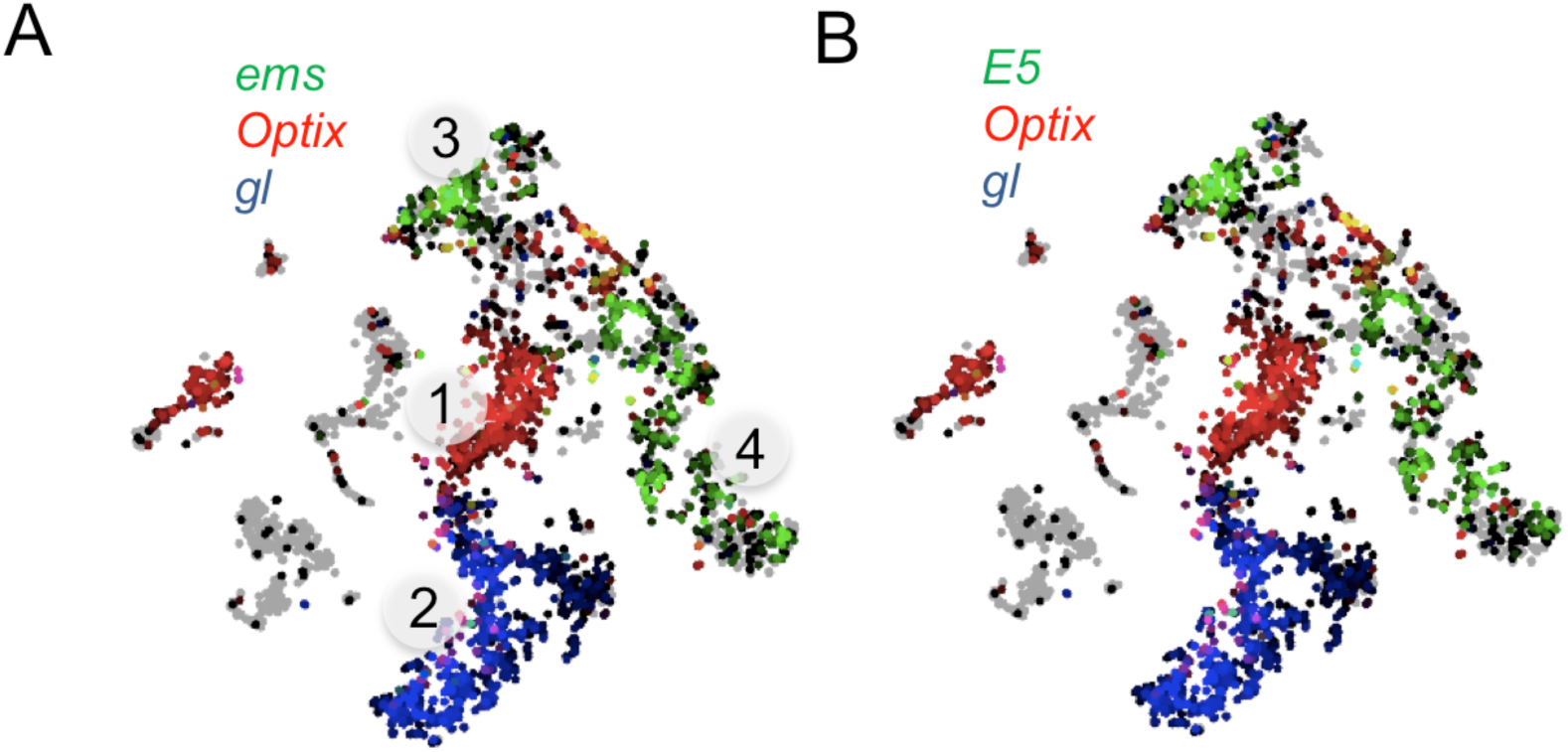
SCOPE visualization of the single cell expression of the paralogous genes *ems* (A) and *E5* (B). The genes *Optix* and *gl* are included to mark the eye progenitor cells (1) and the differentiating retina (2), respectively. *ems* and *E5* are expressed in the prospective head (3) and antenna (4), which are also marked on the cell atlas.

**Figure S4.**
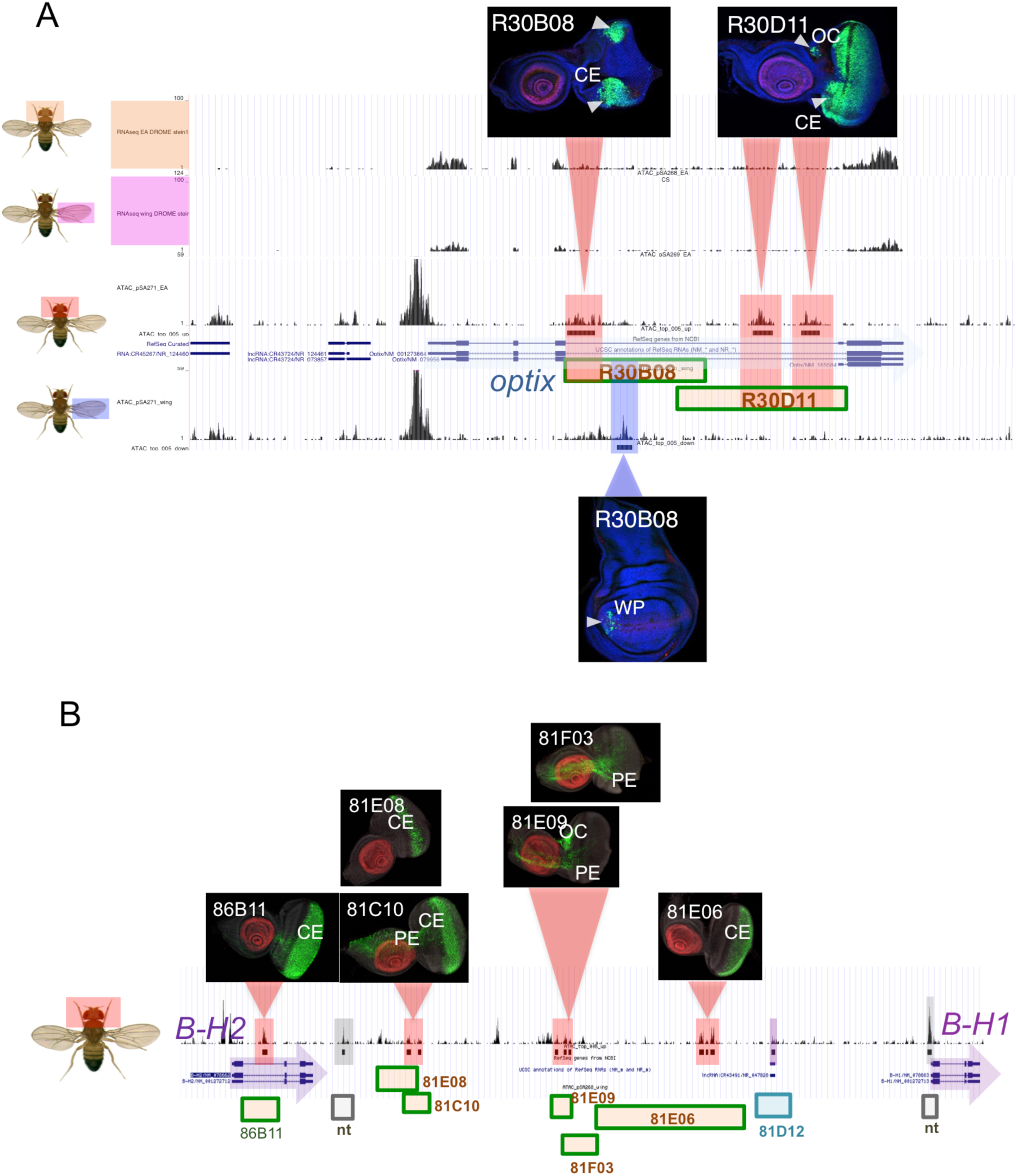
Genome browser view of the *Drosophila Optix* (A) and (B) B-H2/B-H1 loci. (A).First two tracks, eye disc and wing disc transcription: Below, eye disc and wing disc ATACseq tracks. DARs are highlighted. Enhancer activity of the two Flylight lines indicated, as marked with GFP (green; marked by the arrowheads). Above, eye discs; below, wing disc. The discs are also stained for Dll (red) and counterstained with the nuclear marker DAPI (blue). (B) B-H2/B-H1 loci. Eye ATACseq track from genome browser. DARs are highlighted. Above, Flylight images of enhancer activity driven by different regions (green bars). Of the 12 DARs detected, 10 fall within tested regions. Except for 81D12 (highlighted in magenta), for which no imaginal disc enhancer activity is reported (and for which the ATAC peak might derived from the fact that it coincides with an annotated long non-coding RNA, lncRNA:CR43491/NR_047828), all fall within regions with eye disc enhancer activity. Two DARs lie in regions not covered by the Flylight fragments (highlighted in grey). As in (A), the antennal region of the eye-imaginal disc is marked in red. CE: compound eye; OC: ocellar region; PE: peripodial epithelial layer; WP: wing pouch.

**Figure S5.**
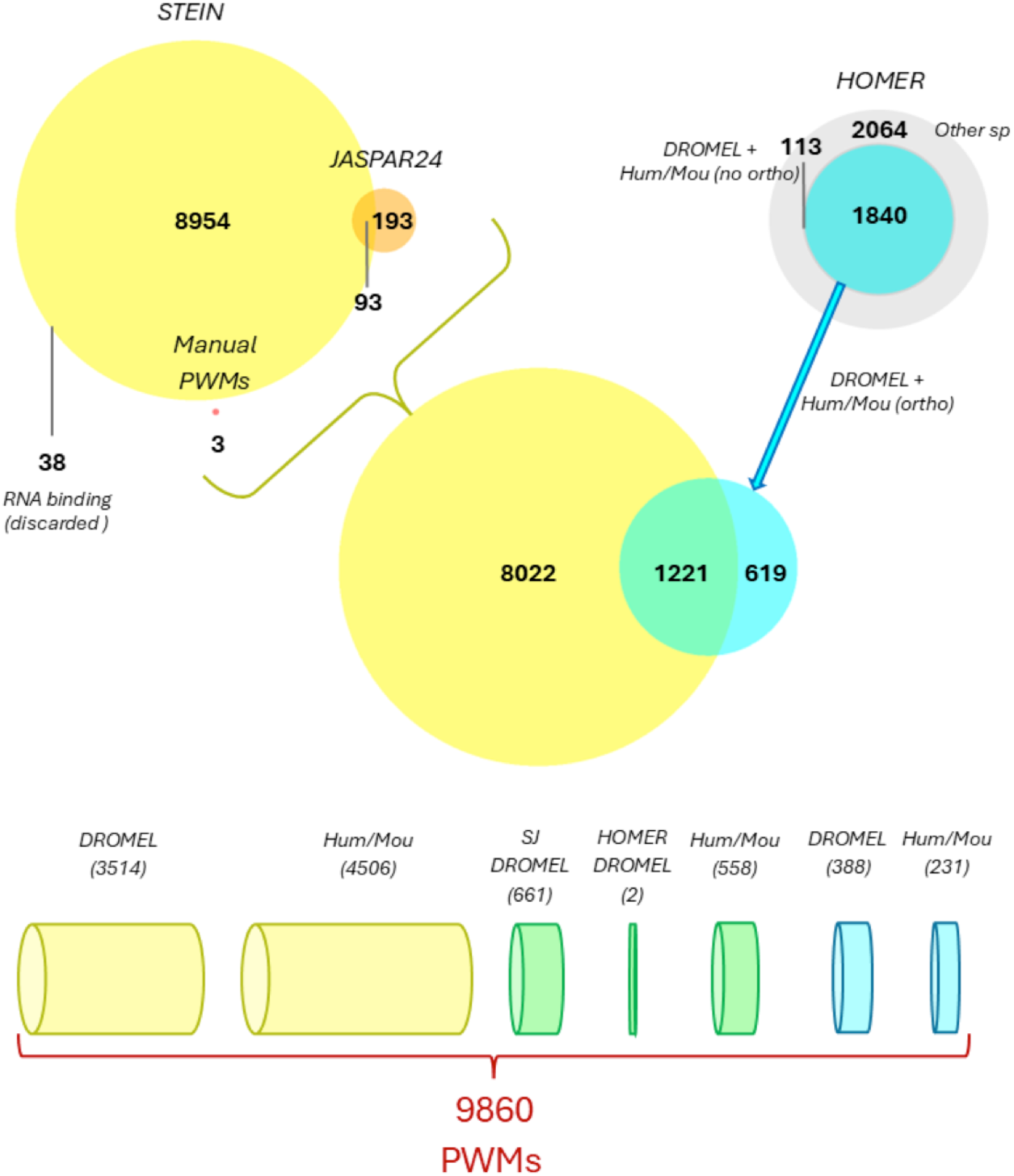
Pipeline for filtering and integrating the motif collections used. The filtered STEIN collection (yellow, top left) is merged with JASPAR24 (orange) and the manually inserted PWMs (red). The HOMER collection is filtered twice (cyan, top right) before calculating the intersection with the other collections. The subsets derived from the intersection (cylinders) are classified according to the origin of the annotations, direct (DROMEL) or indirect (Hum/Mou). The common PWMs (green cylinders) are categorized into direct annotations in the STEIN + JASPAR24 collection (SJ DROMEL), direct annotations in the HOMER collection (HOMER DROMEL), and indirect annotations in the STEIN + JASPAR24 collection (Hum/Mou).

**Figure S6.**
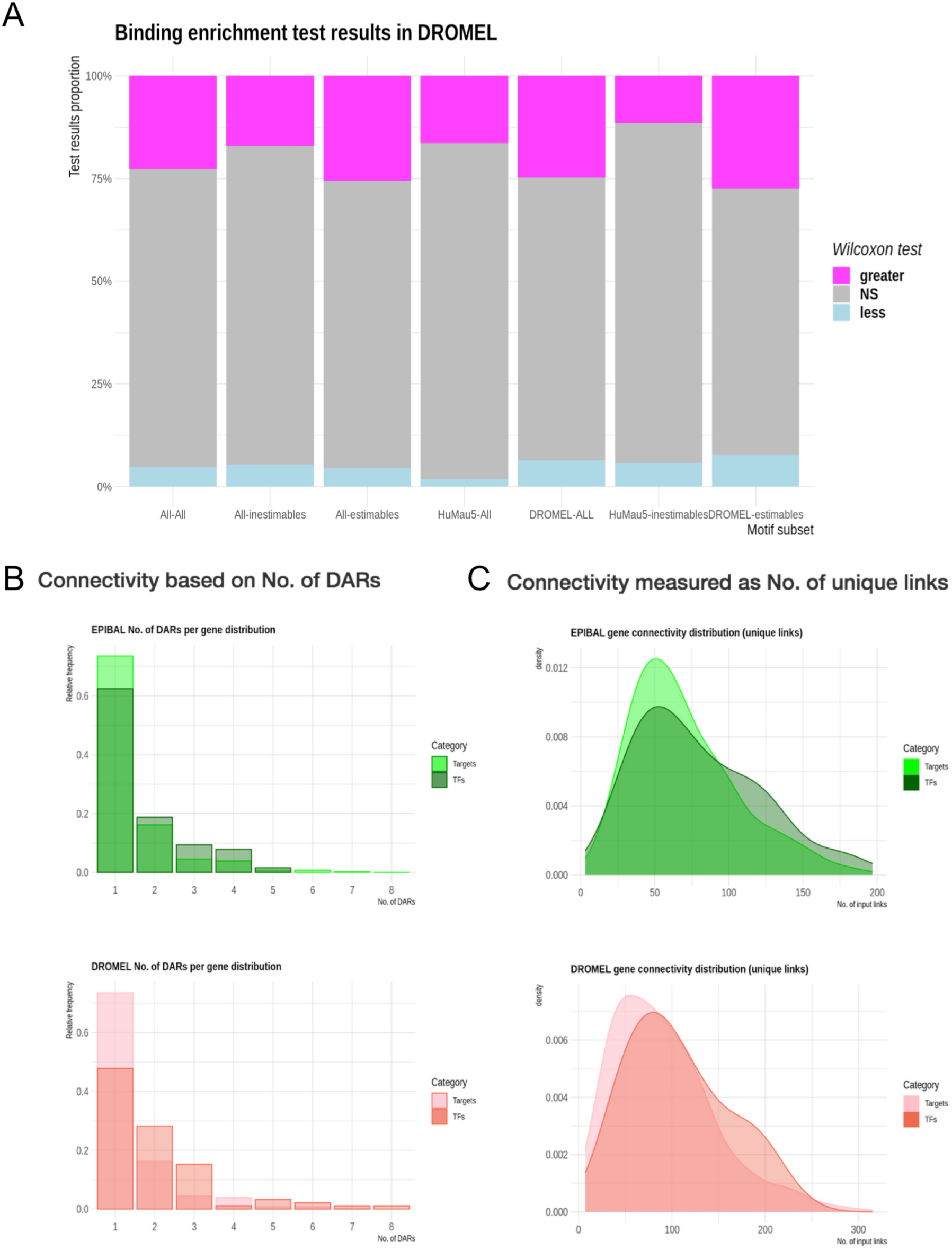
(A) TF motif enrichment in *Drosophila* DARs of eye versus wing, for different subsets of motifs. *Drosophila* estimable motifs were most enriched. (B) The number of eye DARs linked to TF is larger than the number associated to non-TF genes (“targets”) both in *Episyrphus (EPIBAL)* and *Drosophila (DROMEL).* (C) Similarly, the number of unique TF predicted to bind to an eye gene (within the sum of all its DARs) is also larger for TFs than for other genes in both species.

**Figure S7.**
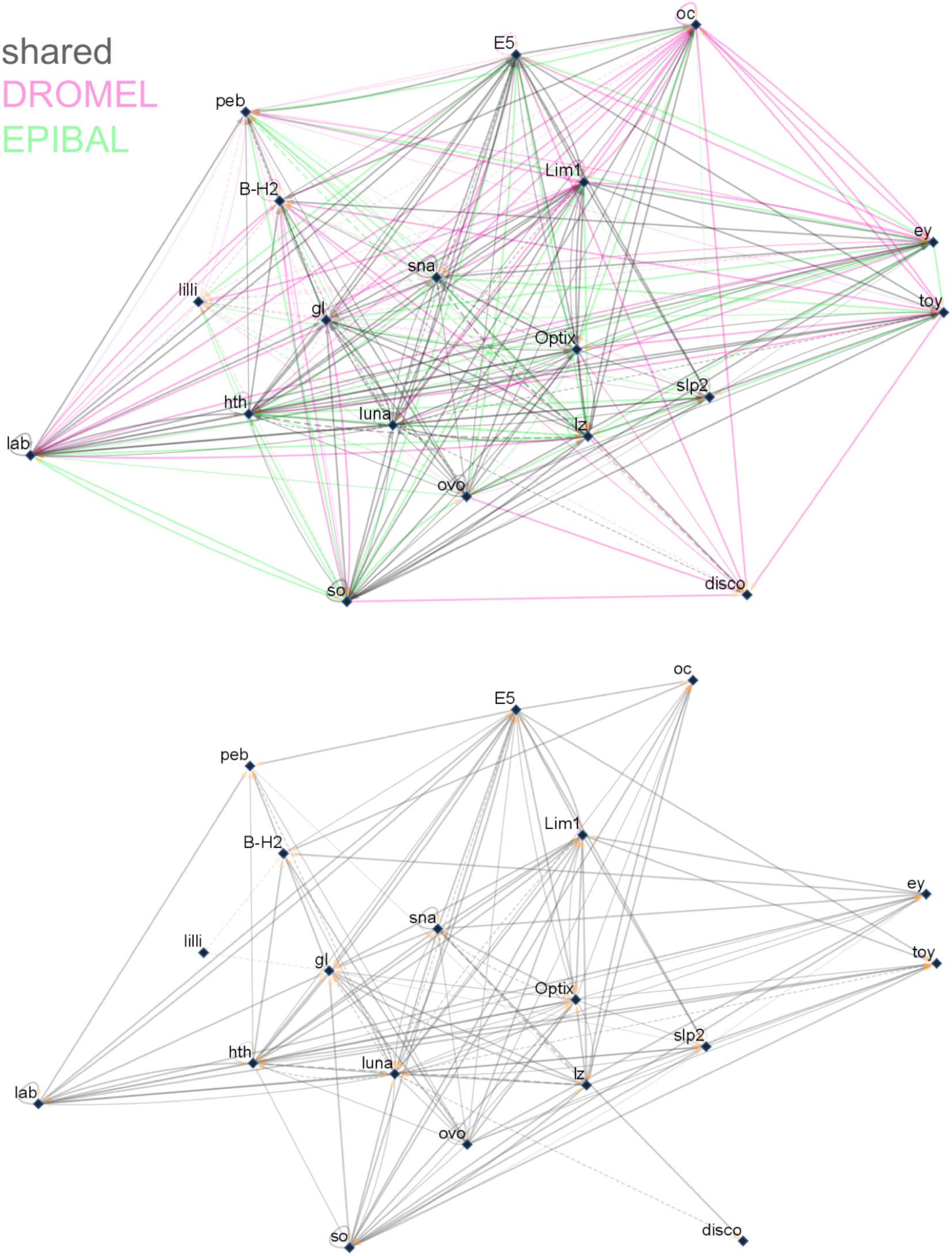
Cytoscape view of the kernel TF GRN. Links are directed, with yellow arrowheads pointing to the regulated targets. DROMEL-specific links, pink; EPIBAL-specific links, green; Shared links, grey. Line type: *solid*: DROMEL motifs; *dashed*: human and mouse motifs; Increasing Line *Transparency*: estimable to non estimable; Increasing Line *Width*: non enriched to enriched in eye discs. The full network (top) and the shared links-only network (bottom) are shown.

**Figure S8 (Supplementary to Figure 3).**
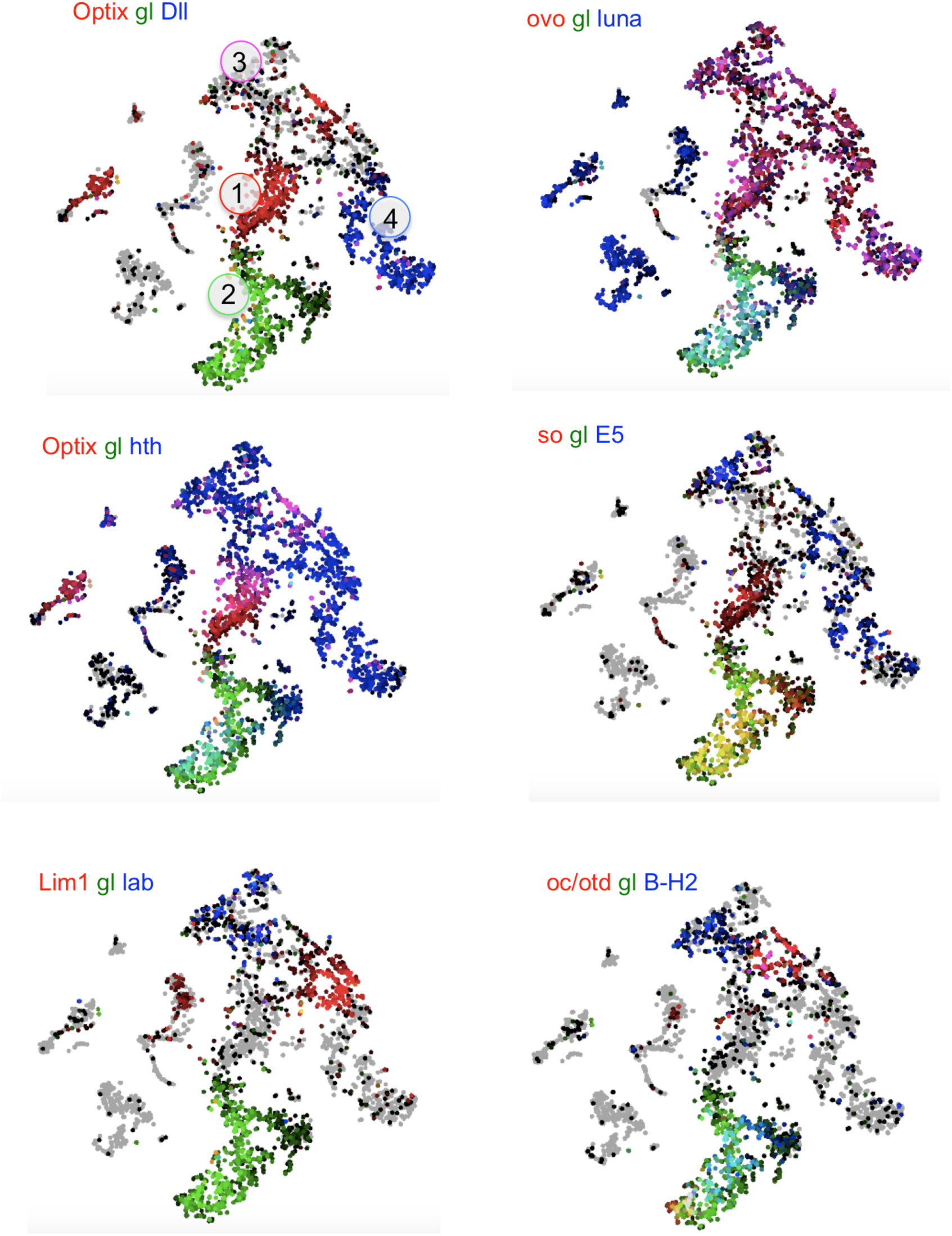
SCOPE visualization of single cell mRNA abundance. (1), (2), (3) and (4) indicate eye progenitors/precursors, differentiating retina, prospective head capsule and antenna, respectively. *gl* expression is represented in green. green/red overlap: yellow. green/blue overlap: cyan. red/blue overlap: magenta. red/green/blue overlap: white.

**Figure S9.**
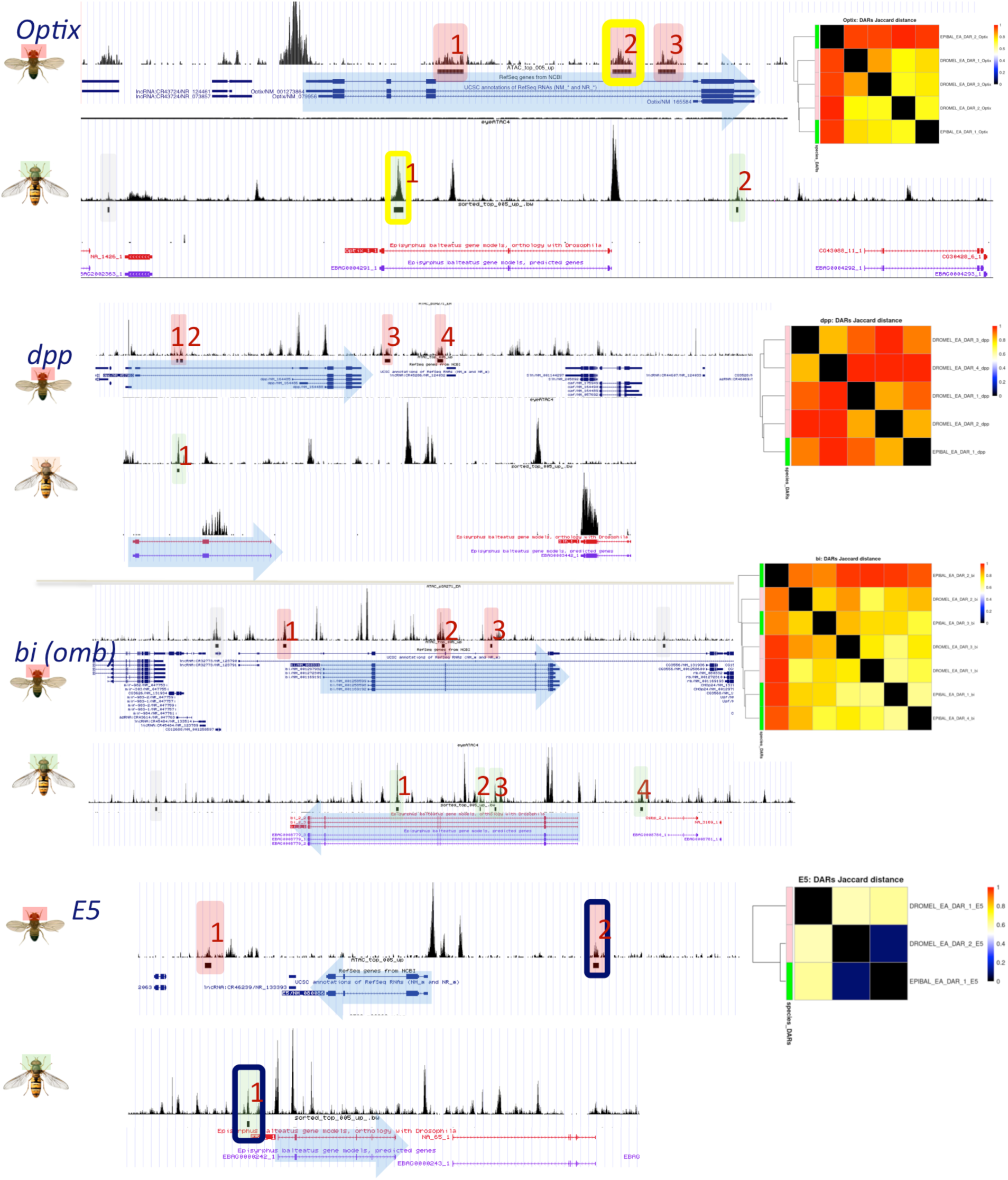
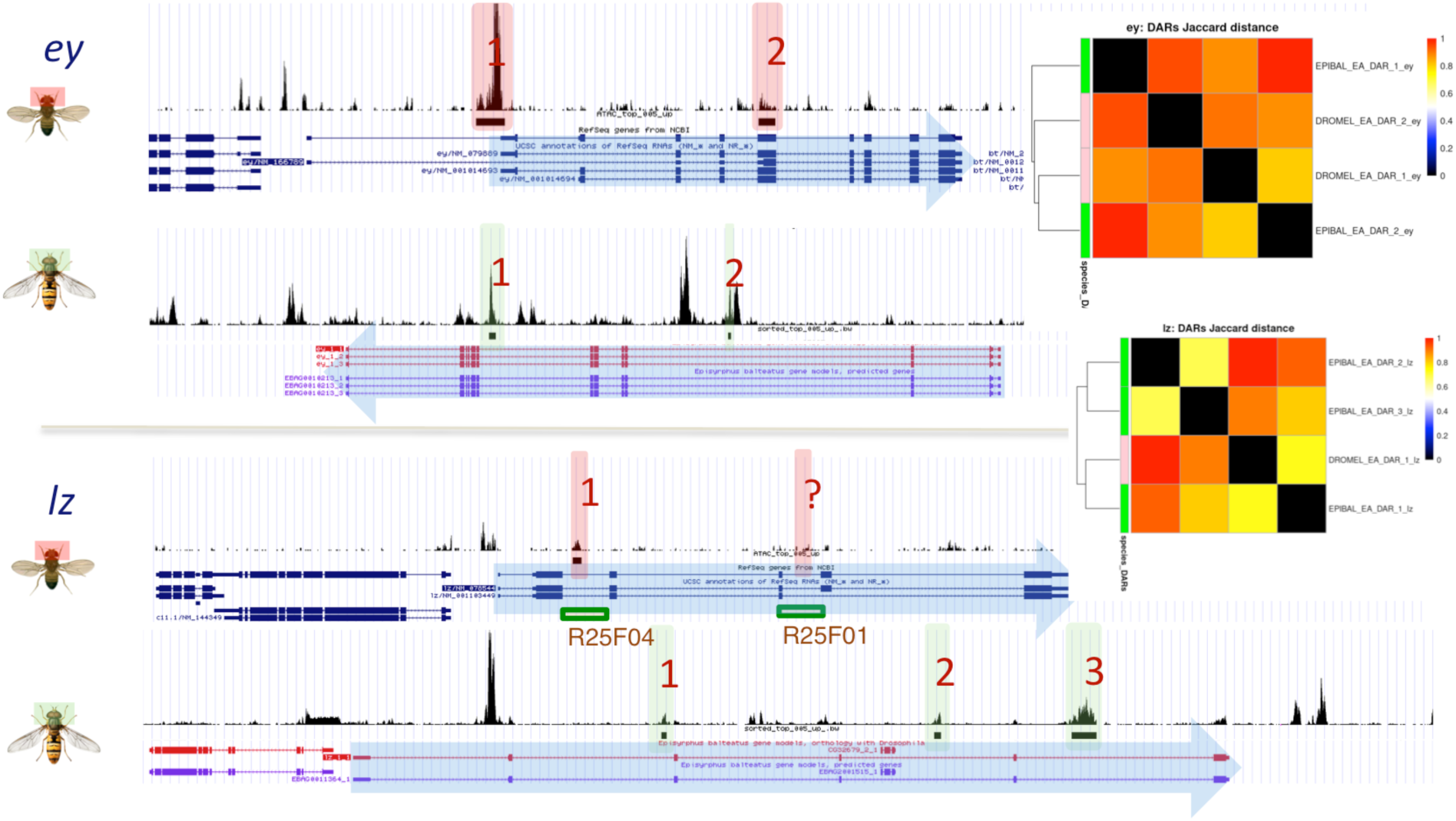
A sample of paralogous loci indicating the number, location and similarity of linked DARs. In the case of *lz,* the Flylight project identified two eye enhancers fragments (boxes) in the *Drosophila* locus. Our ATACseq data seems to have missed one of them (marked as “?”).

**Figure S10.**
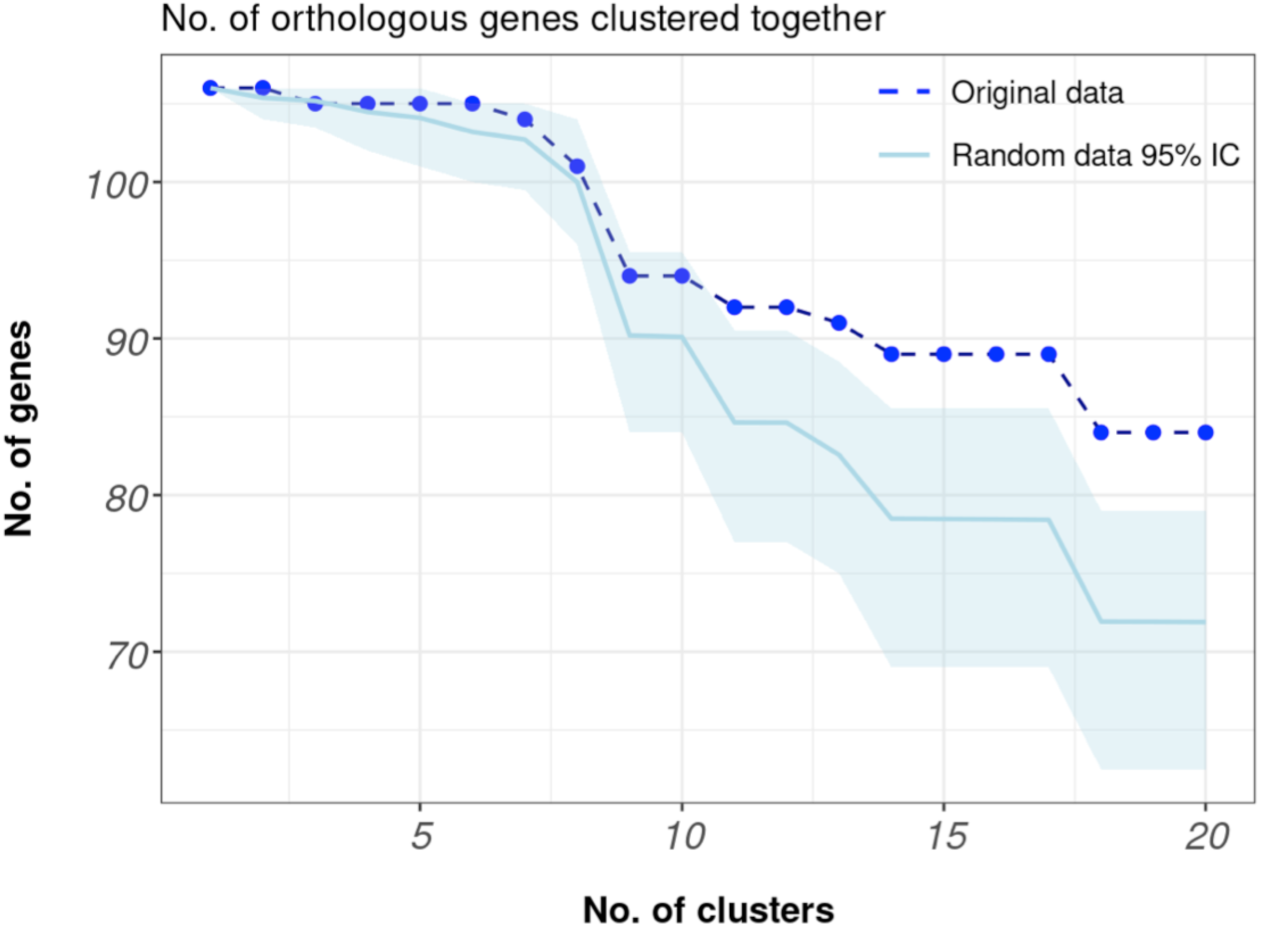
Conservation curve of the regulatory landscape between orthologous gene pairs. The empirical curve (dotted line) represents the decrease in the number of orthologous pairs clustered together (hierarchical clustering) as a function of the number of selected clusters. The gene set was clustered based on the distances between genes (Jaccard distances), calculated from the number of distinct motifs in the kernel of 20 TFs found in any of the DARs that make up the regulatory landscape of each gene. To establish the random distribution of conservation curves (shaded area), each gene was randomly assigned the regulatory landscape of another gene prior to clustering. This process was repeated 100 times, selecting the 95 curves (95% confidence) closest to the mean random curve (solid line). The gene core network shows a higher degree of conservation of regulation across species (i.e. the original data curve lies above the confidence area) than would be expected by chance.

## Supplementary Material

**Supplementary Table S5.**
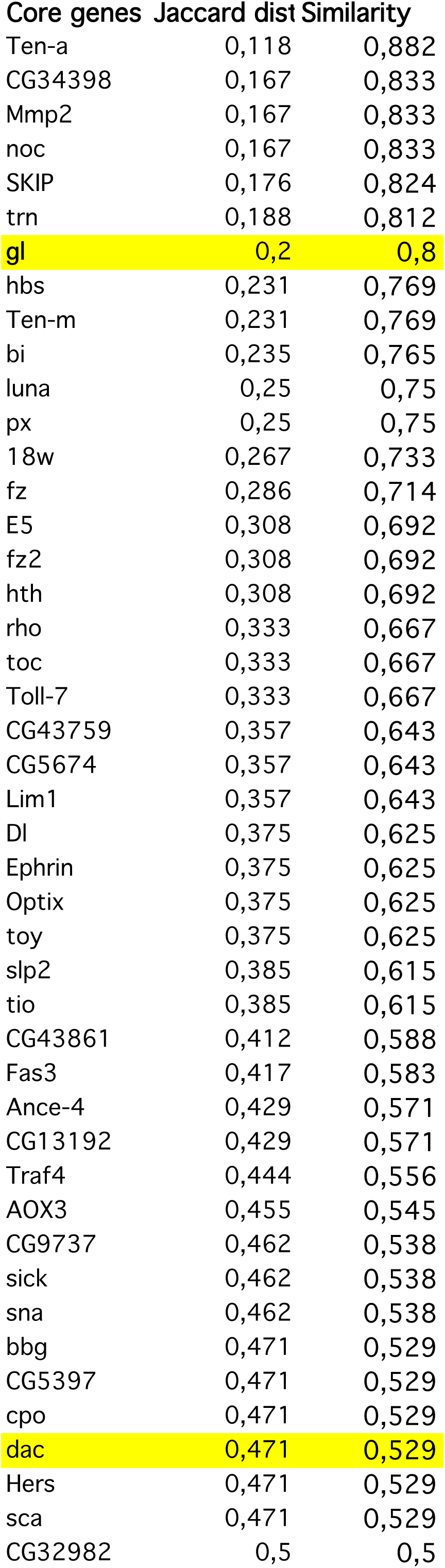
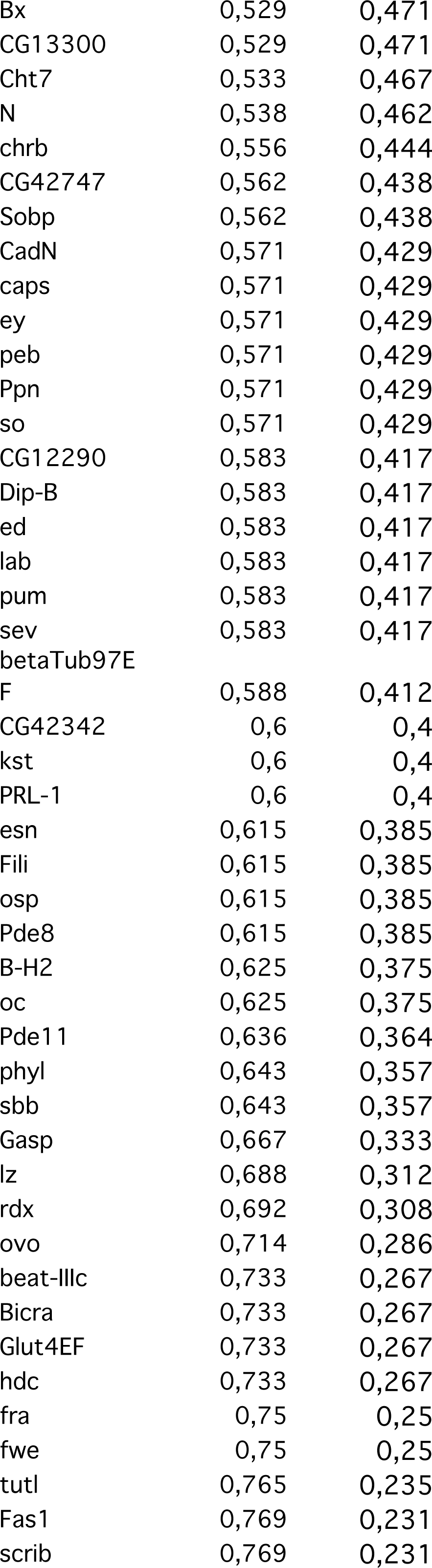
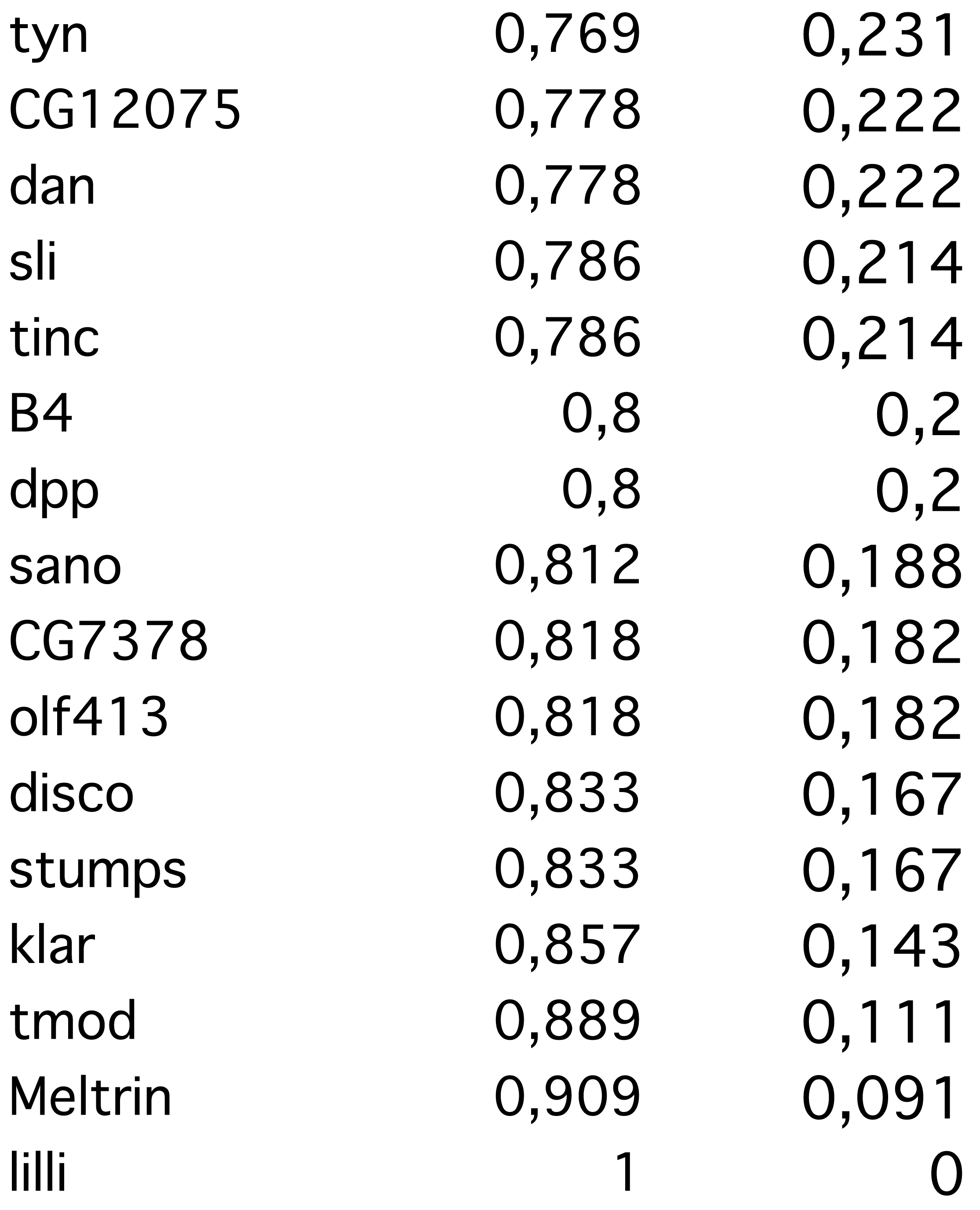
List of 106 core genes. Jaccard distance and Regulatory Similarity index are indicated for each of them.

**Supplementary Table S6.**
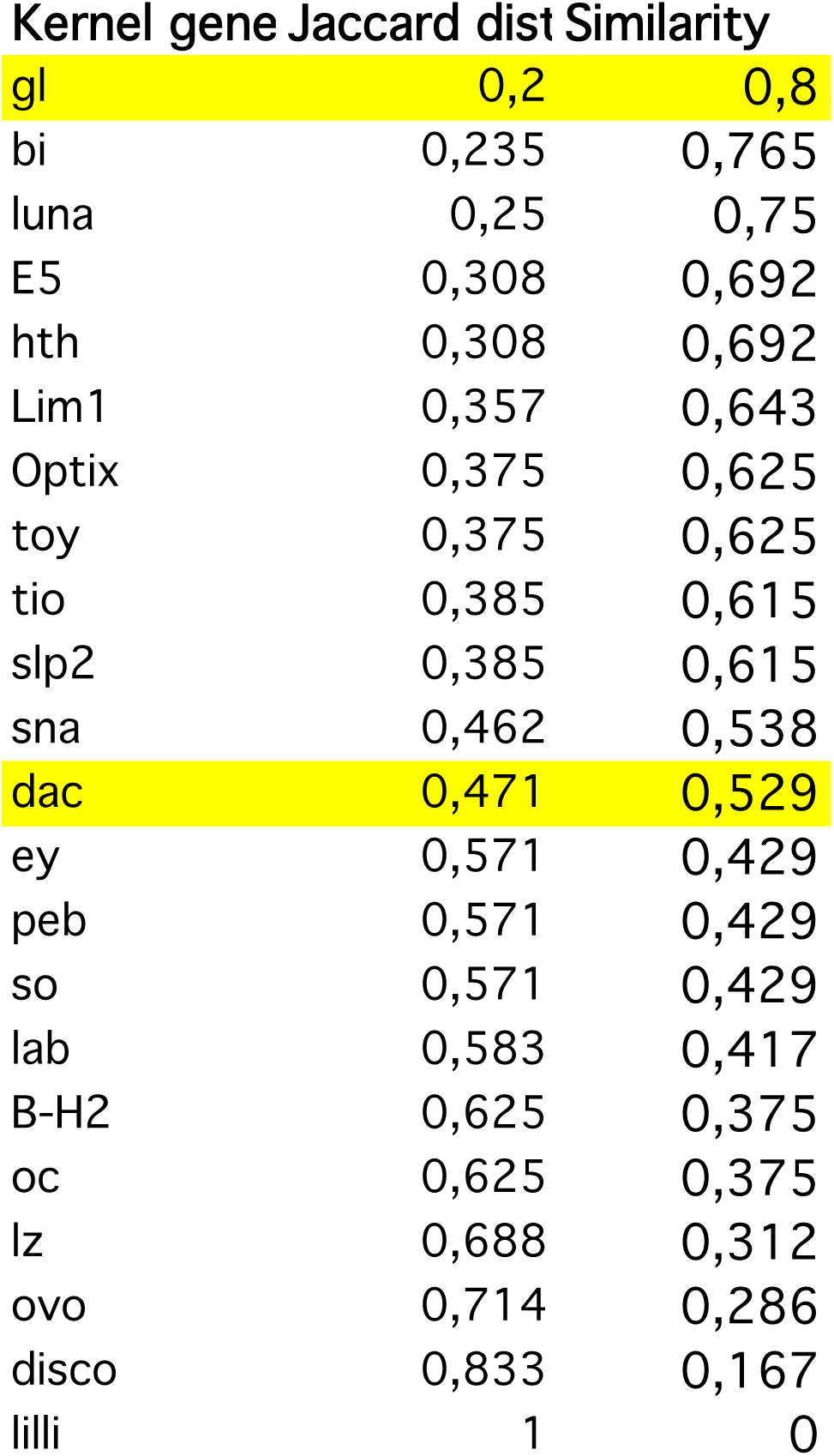
List of 22 kernel transcription factors. Jaccard distance and Regulatory Similarity index are indicated for each of them.

